# ERH elicits cell lineage restriction in mammalian preimplantation development and differentiation from pluripotency via H3K9me3-heterochromatin

**DOI:** 10.1101/2024.06.06.597604

**Authors:** Andrew Katznelson, Blake Hernandez, Kylea Tapia, Holly Fahning, Adam Burton, Jingchao Zhang, Maria-Elena Torres-Padilla, Nicolas Plachta, Kenneth S. Zaret, Ryan L. McCarthy

**Affiliations:** Institute for Regenerative Medicine and Department of Cell and Developmental Biology, Perelman School of Medicine, University of Pennsylvania, Philadelphia, PA 19104, USA; Center for Developmental Biology and Regenerative Medicine, Seattle Children’s Research Institute, Seattle, WA 98101, USA; Institute of Epigenetics and Stem Cells (IES), Helmholtz Zentrum München D-81377 München, Germany; Faculty of Biology, Ludwig-Maximilians Universität, München, Germany; Department of Pediatrics, University of Washington School of Medicine, Seattle, WA 98195, USA

**Keywords:** ERH, H3K9me3, heterochromatin, embryo, pre-implantation, differentiation, pluripotency, naïve

## Abstract

Enhancer of Rudimentary Homolog (ERH) is an evolutionarily conserved protein originally characterized as promoting fission yeast heterochromatin^1^ and recently shown to maintain H3K9me3 heterochromatin in human fibroblasts^2^. Here, we find that ERH depletion in fibroblasts reverts the somatic cell H3K9me3 landscape of broad megabase size domains to an embryonic stem cell (ESC) state composed of mainly H3K9me3 peaks and enables activation of naïve and pluripotency genes and transposable elements during induced pluripotent stem cell (iPSC) reprogramming. Concordantly, we find that ERH represses totipotent and alternative lineage programs during mouse preimplantation development and is required for proper segregation of the inner cell mass and trophectoderm cell lineages. During human ESC differentiation into germ layer lineages, ERH silences naïve and pluripotency genes, transposable elements, and alternative lineage somatic genes. As in fission yeast, we find that mammalian ERH interacts with RNA-binding proteins to engage and repress its chromatin targets. Our findings reveal a conserved, fundamental role for ERH in cell fate specification via the initiation and maintenance of early developmental gene repression.

## Main text

Heterochromatin marked by histone H3 lysine 9 trimethylation (H3K9me3) represses the expression of many protein-coding genes and DNA repeat elements^3–5^. Yet despite its stability, H3K9me3-heterochromatin is dynamically established, maintained, and rearranged during embryogenesis^3,6,7^. Zygotic genome activation (ZGA), marked by the initial transcription of key genes and transposable elements^8^, occurs at the 1-2 cell stage in mice and the 4-8 cell stage in humans^8^ and requires demethylation of maternal H3K9me3 marks created in oocytes^9,10^. As the embryo loses totipotency and differentiates into the first two developmentally-specified lineages, the trophectoderm of the blastocyst and the inner cell mass (ICM), many ZGA-specific protein-coding genes and repeats gain H3K9me3 and are subsequently silenced ^6,11,12^. As development proceeds, H3K9me3 is further established on lineage-inappropriate genes and repeat elements ^6,11,13^, with the highest number of genes marked by H3K9me3 at the germ-layer stage^3^. Throughout germ layer specification to mature somatic lineages, H3K9me3 is removed from genes that become active and is deposited onto early developmental genes that become repressed^3^. The H3K9me3 landscape changes substantially from pluripotency to early and terminally differentiated states^14,15^. The mechanisms operating in the initial targeting of histone methyltransferases to deposit H3K9me3 and the maintenance of the initially established H3K9me3 domains remain poorly understood. In this study, we compared how *enhancer of rudimentary homolog* (ERH), a highly conserved heterochromatin protein distinct from histone methyltransferases, contributes to H3K9me3 dynamics and gene repression in differentiated somatic cells, pluripotent cell lines, and early mammalian development.

ERH is an evolutionarily conserved protein originally discovered in fission yeast to regulate sporulation, the primary form of yeast cell differentiation, via H3K9 methylation ^1,16^. In *S. pombe,* Erh1 forms a complex with the RNA binding protein Mmi1, which is recruited by binding of nascent meiotic transcripts through its YTH domain^1,15^. Erh1-Mmi1 then causes the degradation of its bound RNA transcripts, the establishment of H3K9 methylation, and consequent repression of meiotic and developmental genes, transposons, and rDNA^1,16^. We recently found that in terminally-differentiated primary human fibroblasts, ERH is highly enriched in H3K9me3-heterochromatin, where it represses meiotic and germ cell genes, similar to that observed in fission yeast^1,2,5^. We further found that ERH in mammals has evolutionarily expanded its function to include the majority of regions that exhibit H3K9me3 heterochromatin in somatic cells, including alternative-lineage protein coding genes and repeat elements^2^. While ERH is highly conserved among eukaryotes, its fission yeast partner Mmi1 lacks a metazoan homolog^17^. Thus, how cell-type specific cofactors recruit ERH to chromatin to regulate cell identity throughout mammalian development is unknown.

Here, we find that upon ERH loss in fibroblasts, there is reversion to a pluripotency-like H3K9me3 state which is permissive to binding of pluripotency transcription factors, which in turn activate pluripotency and alternative lineage genes and repeat elements. The derepression of pluripotency and alternative lineage programs led us to investigate the role of ERH in the earliest developmental stages, where repression by H3K9me3 has been implicated during trophectoderm and inner cell mass segregation in the blastocyst^6,11,12^ and the differentiation from pluripotency to the germ layer lineage^3^. By genetic perturbation, chromatin and gene expression profiling, and imaging, we identify a role for ERH across mouse and human systems in repressing early naïve pluripotency and alternative lineage genes. We find that targeting of mammalian ERH to chromatin occurs through cell-type specific interactions with RNA binding proteins, enabling context-dependent functions. Together, our findings demonstrate a fundamental role for ERH during developmentally acquired H3K9me3-mediated gene repression and maintenance of such repression in somatic cells.

## Results

### ERH depletion in primary human fibroblasts causes H3K9me3 to revert to an ESC-like state

We assessed how H3K9me3 is disrupted after ERH loss by CUT&RUN for H3K9me3 after siRNA knockdown in human primary fibroblasts (Fig. 1a). Consistent with our previous study^2^, we confirmed that H3K9me3 is globally reduced in human fibroblasts after ERH knockdown (Fig. 1b). Yet while large megabase-scale domains were diminished (Fig. 1c top, purple domains in top two tracks), as seen previously^2^,10,526 H3K9me3 residual peaks were retained in the ERH knockdown (Fig. 1c, lower, second track) and Supplementary Table 1). Remarkably, the residual H3K9me3 peaks in siERH fibroblasts resemble the H3K9me3 profile in H1 ESCs (Fig. 1c lower, third track). Across the genome, 85% of the residual fibroblast siERH H3K9me3 sites overlap with H3K9me3 peaks in H1 hESCs (Extended Data Fig. 1a). Pearson correlation of H3K9me3 peaks from fibroblasts after siControl or siERH, as well as in hESCs, supports a shift in the H3K9me3 heterochromatin landscape to an embryonic state (Fig. 1d,e). Thus, while loss of ERH in primary fibroblasts causes a global loss of H3K9me3 domains, embryonic H3K9me3-marked targets are retained.

**Figure 1.**
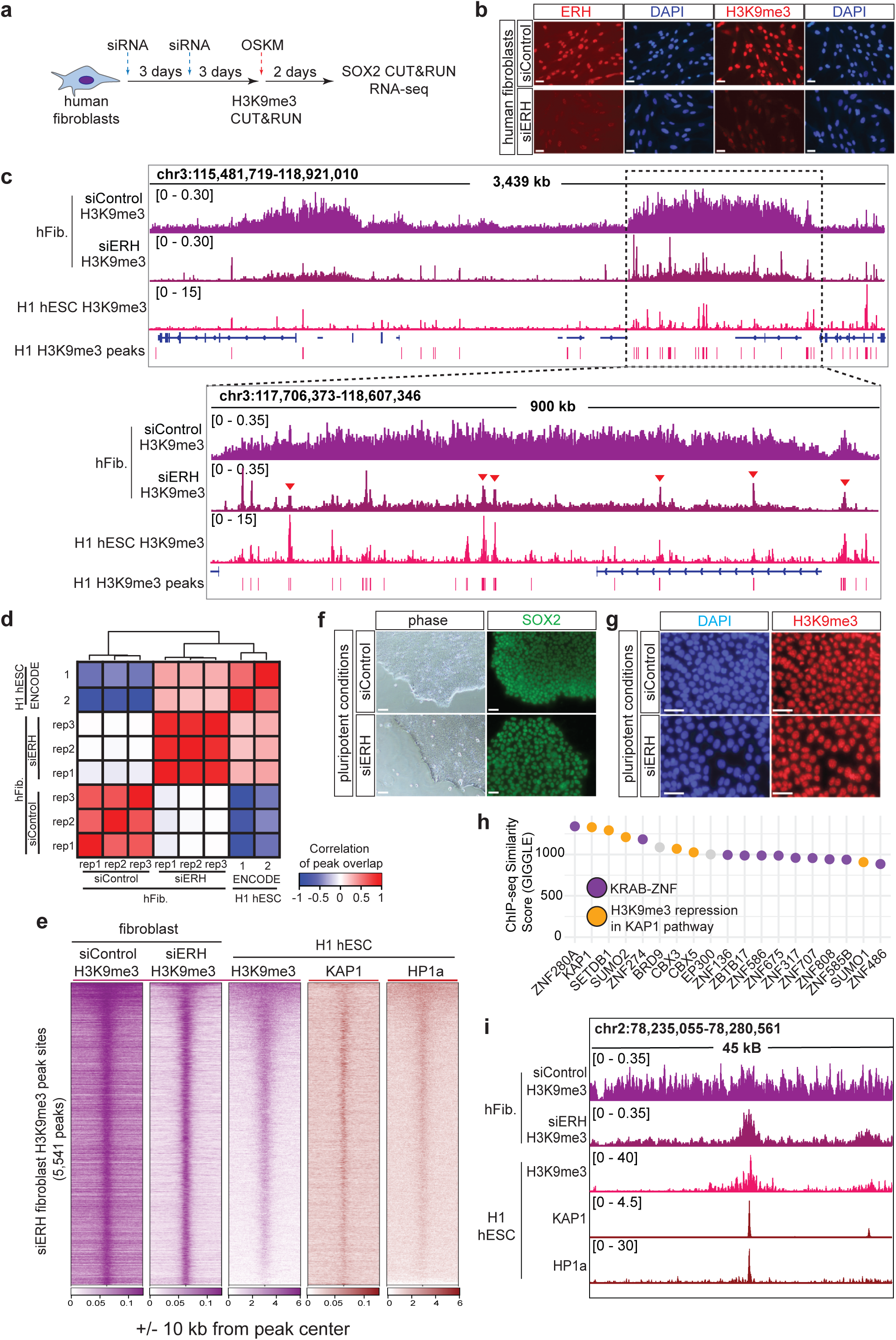
ERH depletion reverts fibroblast H3K9me3 patterns to an hESC-like state. **a.** Schematic of siRNA treatments and timing for H3K9me3 CUT&RUN experiments shown in figure 1 and SOX2 CUT&RUN and RNA-seq experiments shown in figure 2. **b.** Immunofluorescence of ERH (red), H3K9me3 (red) and DAPI (blue) in siControl and siERH human fibroblasts. Scale bars, 50µm. **c.** Browser track of H3K9me3 levels in siControl and siERH treated human fibroblasts and H1 hESCs (GSM1003585) at the megabase-domain level and gene level. Example H3K9me3 peaks common between siERH fibroblasts and H1 hESCs are indicated by red triangles. **d.** Heatmap of fibroblast siControl and siERH H3K9me3, H1 hESC H3K9me3 (GSM1003585) and H1 hESC HP1α (GSE123229) levels centered at siERH retained H3K9me3 peaks located within fibroblast H3K9me3 domains +/-10kb. **e.** Correlation of H3K9me3 from BJ fibroblasts after siControl or siERH with H1 hESC H3K9me3. **f.** Representative images showing hESC morphology and SOX2 (green) pluripotency marker staining in siControl and siERH H1 hESCs under pluripotency maintaining conditions. Scale bars, 50µm. **g.** Representative images of H3K9me3 (red) and DAPI (blue) in siControl and siERH H1 hESCs under pluripotency maintaining conditions. Scale bars, 50µm. **h.** CistromeDB Toolkit enrichment analysis of fibroblast siERH H3K9me3 peaks. The top 20 proteins scored by the analysis are visualized. Proteins colored by characterization as H3K9me3-associated (orange), KRAB zinc-finger transcription factors (purple), and euchromatin-associated (gray). **i.** Browser track comparing siControl and siERH fibroblast H3K9me3 with H1 hESC H3K9me3 (GSM1003585) and HP1a (GSM1701825) patterns. Peak locations called by SEACR analysis on siERH H3K9me3 data are shown. See also Extended Data Fig. 1 and Supplementary Table 1.

We hypothesized that since ERH depleted fibroblasts retain hESC-like H3K9me3 peaks (Fig. 1c-e), ERH may be dispensable for maintaining H3K9me3 in hESCs. To test this hypothesis we depleted ERH in H1 hESCs cultured in pluripotency-maintaining conditions, which caused no observable changes in colony morphology or loss of pluripotency (Fig. 1f), even after sustained knockdown (Extended Data Fig. 1d). H3K9me3 level and nuclear organization by immunofluorescence was not changed in siERH-treated hESCs (Fig. 1g; Extended Data Fig. 1e). We also observed no change in H3K27me3 (Extended Data Fig. 1f). Despite no apparent disruption of the pluripotent state or global H3K9me3 levels, ERH-depleted hESCs show an increased nuclear area and nuclear blebbing (Extended Data Fig. 1g), which are changes associated with chromatin decompaction^18^. We previously found that ERH depletion did not change the expression or protein level of H3K9me3 HMTs^2^, we similarly observe no significant decrease in H3K9me3 HMT expression upon ERH depletion in pluripotent hESCs (Extended Data Fig. 1h). To investigate potential locus-specific changes in hESC H3K9me3 upon ERH depletion, we conducted H3K9me3 CUT&RUN in siControl and siERH treated hESCs. Consistent with our immunofluorescence results, we observed minimal change in H3K9me3 (Extended Data Fig. 1i) and high correlation between H3K9me3 peaks in siControl and siERH treated hESCs (Extended Data Fig. 1j).

To identify chromatin-associated proteins and pathways regulating the residual H3K9me3 peak sites in human fibroblasts, we analyzed the regions using the cistromeDB Toolkit, which uses a compendium of published ChIP-seq data sets^16,17^. We find that most proteins enriched at the residual peak sites in ES cells (i.e., prior to ERH knockdown) are either KRAB-ZNFs or involved in H3K9me3 repression via KAP1, which is recruited to chromatin through binding the KRAB domain (Fig. 1h)^19^. Likewise, de novo motif analysis finds that enriched sequence motifs in the residual peak sites are for KRAB-ZNFs factors (Extended Data Fig. 1k). Consistent with the enrichment analysis, the residual siERH H3K9me3 peak sites are highly enriched for H3K9me3, KAP1, and HP1 in ES cells (Fig. 1e,i; Extended Data Fig. 1l). As KRAB-ZNFs repress transposons via H3K9me3, we hypothesized the residual peaks would be enriched over repeats. Repeat enrichment analysis found that the residual peaks are enriched over canonical KRAB-ZNF targets: LINEs, LTRs, and satellite repeats (Extended Data Fig. 1m). We conclude that ERH is required to maintain the broad H3K9me3 domains that are established over the course of development and are present in terminally differentiated cells, but does not control H3K9me3 peaks attributed to the DNA-dependent association of KAP1 that persists from pluripotency over constitutively repressed repeats.

### ERH-dependent H3K9me3 suppresses SOX2 binding and activation of pluripotency genes and repeats during OSKM reprogramming

H3K9me3 impairs induced pluripotent stem cell (iPSC) reprogramming by blocking transcription factor binding, and pluripotency genes marked by H3K9me3 show delayed activation compared to genes in heterochromatin marked with H3K27me3 or neither mark (Extended Data Fig. 2a)^20^. We therefore assessed the impact of ERH depletion in the context of OSKM transcription factor binding and gene activation during iPSC reprogramming. We focused on a 48 hour time point after initiation of iPSC reprogramming (Fig. 1a and Supplementary Tables 2-4), as H3K9me3 is inhibitory to transcription factor activity at this stage^20^.

CUT&RUN of SOX2 after 48 hours of OSKM reprogramming showed that while most binding sites were shared between siControl and siERH fibroblasts, 2,538 binding sites were gained (Extended Data Fig. 3a and Supplementary Table 2). We observed that ERH knockdown enabled SOX2 to target loci that lose H3K9me3, including within heterochromatin domains previously annotated as resistant to reprogramming factor binding in fibroblasts^20^ (Fig. 2a, and Extended Data Fig. 3b-d). By assessing the genomic locations of siERH-gained SOX2 peaks in the cistromeDB toolkit^21,22^, we found that the newly gained SOX2 sites had the most similarity to ChIP-seq peak sets of SOX2 and its pluripotency partners NANOG and OCT4 in embryonic stem cells (Extended Data Fig. 3e). Inspection of genome browser tracks identified specific loci where loss of ERH-controlled H3K9me3 enabled SOX2 binding over normal SOX2 and NANOG targets in ES cells (Fig. 2b, Extended Data Fig. 3f). We next assessed if loss of ERH-dependent H3K9me3 enables other transcription factors to bind previously refractory sites by performing CUT&RUN footprinting of SOX2’s primary binding cofactors during reprogramming, KLF4 and OCT4, and find enhanced footprints for each factor at SOX2 sites gained by siERH (Extended Data Fig. 3g). Thus, ERH loss during pluripotency reprogramming enables transcription factor binding in what were heterochromatic H3K9me3 regions.

**Figure 2.**
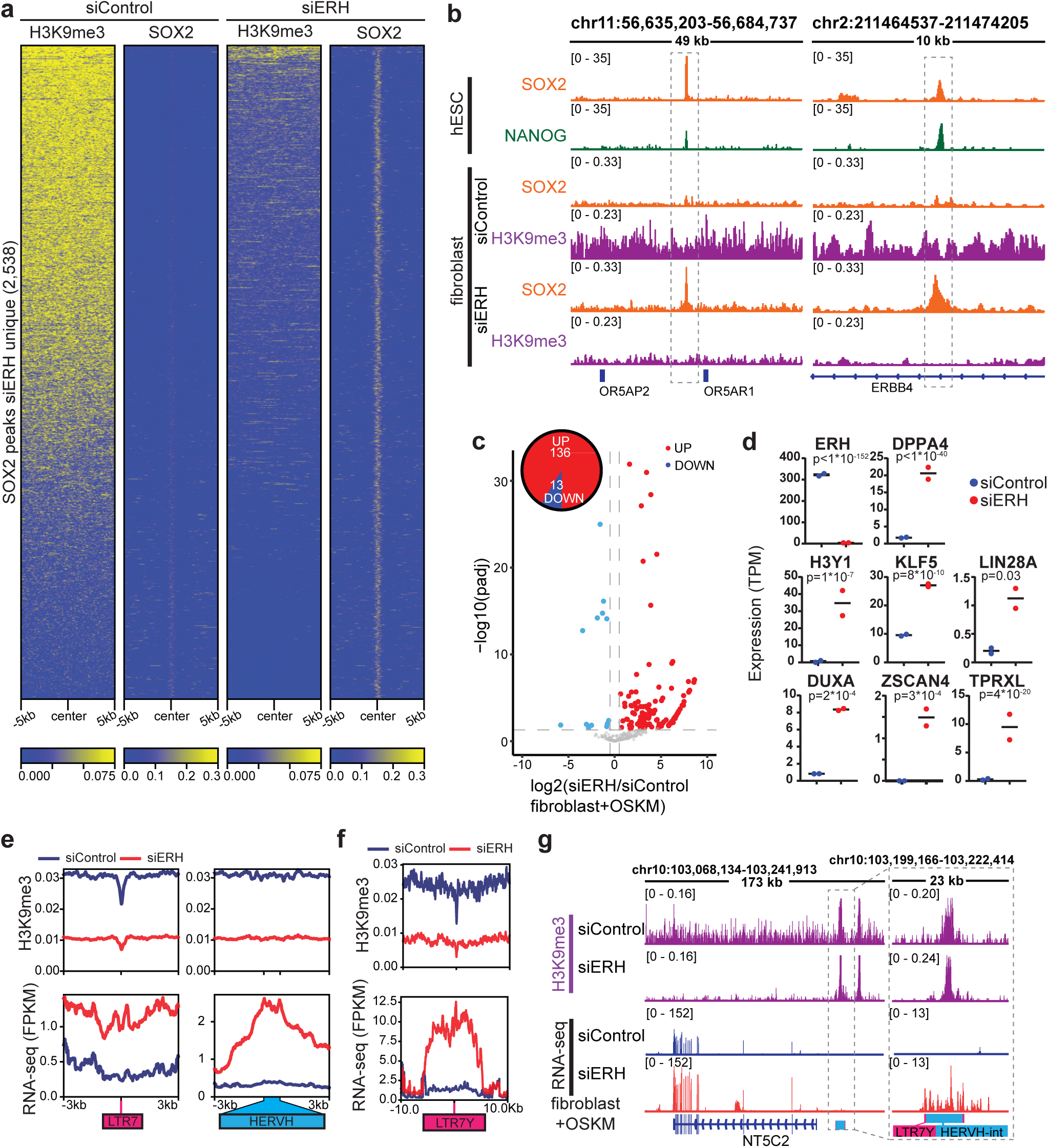
ERH depletion enhances gene activation of pluripotency and alternative lineage genes during iPSC reprogramming. **a.** Heatmap of H3K9me3 CUT&RUN from siControl and siERH human fibroblasts and SOX2 CUT&RUN from siControl and siERH fibroblasts + OSKM at siERH-gained SOX2 peaks. **b.** Browser track of example hESC SOX2 and NANOG peak coincident with fibroblast siERH H3K9me3 loss and SOX2 peak gain. **c.** Volcano plot of expression changes (DESeq2) in siERH vs siControl fibroblasts+OSKM of iPSC activated genes located in fibroblast H3K9me3 domains. **d.** Expression of ERH and example naïve pluripotency genes activated in 48 hours OSKM by ERH depletion, transcripts per million (TPM) values from RNA-seq replicates shown (DESeq2 adjusted p-value, n=2). **e.** Profile plots of H3K9me3 and RNA levels from LTR7 and HERVH repeats in siERH vs siControl fibroblasts+OSKM. **f.** Profile plots of H3K9me3 and RNA levels from LTR7Y repeats in siERH vs siControl fibroblasts+OSKM. **g.** Browser track showing H3K9me3 in siControl and siERH fibroblasts and RNA levels in siControl+OSKM and siERH+OSKM for gene and adjacent LTR7Y naïve pluripotency repeat. See also Extended Data Fig. 2-4 and Supplementary Tables 2-4.

Protein coding genes embedded in H3K9me3 domains, normally refractory to transcription factor activation, were activated after siERH (136 up, 13 down; Fig. 2c). Significantly, genes associated with the human 8-cell like cell (8CLC) state^23^ were activated, including *ZSCAN4, DUXA, DPPA4, LIN28A, TPRXL*^24^*, KLF5,* and the histone variant *H3.Y*^25^ (Fig. 2d and Extended Data Fig. 4a). We compared the activation levels of these genes in siERH+OSKM with existing RNA-seq datasets^26^ and found that *H3.Y, DUXA* and *TPRXL* do not normally get activated at similar levels to siERH+OSKM at any stage during iPSC reprogramming (Extended Data Fig. 4b). However, activation of 8CLC genes in siERH+OSKM is incomplete, as many genes activated in naïve ESCs^27^ are not expressed or expressed at lower levels in siERH+OSKM (Extended Data Fig. 4c), suggesting low activation of naïve programs but not full conversion to a naïve cell state. While the activation of marker genes such as DUXA and ZSCAN4 is lower than in 8C embryos^28^, we conclude that ERH-dependent H3K9me3 safeguards early totipotent and pluripotent genes and that ERH loss creates a permissive environment for transcription factor binding and activation.

We observed that *LTR7* and *HERVH* repeats, which are expressed in and required to maintain pluripotency^29,30^, lose H3K9me3 after ERH depletion and are activated in siERH+OSKM (Fig. 2e). *LTR7Y* repeats, which promote naïve pluripotency^31^, exhibited similar ERH-dependent H3K9me3 and repression (Fig. 2f,g and Extended Data Fig. 4d). Notably, residual H3K9me3 peaks retained at *LTR7Y* repeats were insufficient to maintain repression (Fig. 2f,g). Furthermore, we observed siERH+OSKM activated repeat elements that function specifically during the human 8 cell-like cell stage, and are later marked by H3K9me3 and silenced in the human blastocyst^6^ (Extended Data Fig. 4e). These results demonstrate that ERH maintains H3K9me3 repression of a subset of 8-cell stage and pluripotency repeats in terminally differentiated human cells.

Depletion of H3K9 HMTs has been shown to increase iPSC reprogramming^20^. However, despite increased activation of pluripotency and 8-cell genes at 48 hours after initiation of OSKM-based iPSC reprogramming in siERH treated fibroblasts, by day 2 we observed substantial cell death that persisted 7 days into reprogramming, past the initial apoptotic wave (Extended Data Fig. 4f,g) and resulting in no iPSC colonies forming in siERH+OSKM conditions, compared to the abundant colonies in siControl+OSKM conditions (Extended Data Fig. 4h,i). The failure to generate iPSC colonies was not rescued by simultaneous knockdown of p53 (Extended Data Fig. 4i). Lower ERH siRNA decreased cell death, but did not change the emerging iPSC colony number (Extended Data Fig. 4g,h). Using XDeathDB^32^ on our RNA-seq data identified increased expression of genes activated in several cell death pathways, including RIPK2 signaling (Extended Data Fig. 4j). We investigated if simultaneous inhibition of RIPK2 along with inhibition of RIPK3 and Caspases 3 and 7, which has been shown to improve survival of cells activating 8 cell like genes^23^, could enable survival of siERH+OSKM iPSCs. Notably, we found that adding small molecule inhibitors GSK2983559, Ac-DEVD-CHO, and GSK872 to inhibit RIPK2, Caspases 3 and 7, and RIPK3, respectively, during reprogramming enabled the generation of iPSC colonies with ERH depletion at a significantly higher efficiency compared to siControl (Extended Data Fig. 4k). RIPK2 inhibition alone was necessary and sufficient to successfully generate iPSC colonies expressing pluripotency factors during ERH depletion, while inhibition of RIPK and Caspases 3 and 7 was not sufficient but did increase colony size (Extended Data Fig. 4l). These findings demonstrate that the increased accessibility for transcription factor binding in H3K9me3 domains (Fig. 2a,b) and improved activation of the pluripotency network (Fig. 2c,d) observed upon ERH depletion can increase iPSC reprogramming efficiency.

### ERH is necessary for trophectoderm development, the first embryonic differentiation

Our finding that ERH represses naïve genes that are normally expressed in 2-8 cell embryos, in fully differentiated fibroblasts, led us to investigate if ERH establishes silencing of 2-8 cell genes during pre-implantation development. We first analyzed existing RNA-seq data of mouse embryogenesis timepoints from zygote to blastocyst^33^ and found that ERH mRNA is present across pre-implantation development and is significantly increased from the 8-cell stage through the blastocyst stage (Extended Data Fig. 5a). Immunostaining mouse embryos for ERH protein at the zygote, 2-cell, 8-cell and blastocyst stages revealed that, consistent with its mRNA levels, ERH is expressed and nuclear-localized at all stages analyzed, with increasing abundance in the early and late blastocyst stages (Fig. 3a,b). At the blastocyst stage, ERH levels are similar in ICM and trophectoderm cells (Extended Data Fig. 5b).

**Figure 3.**
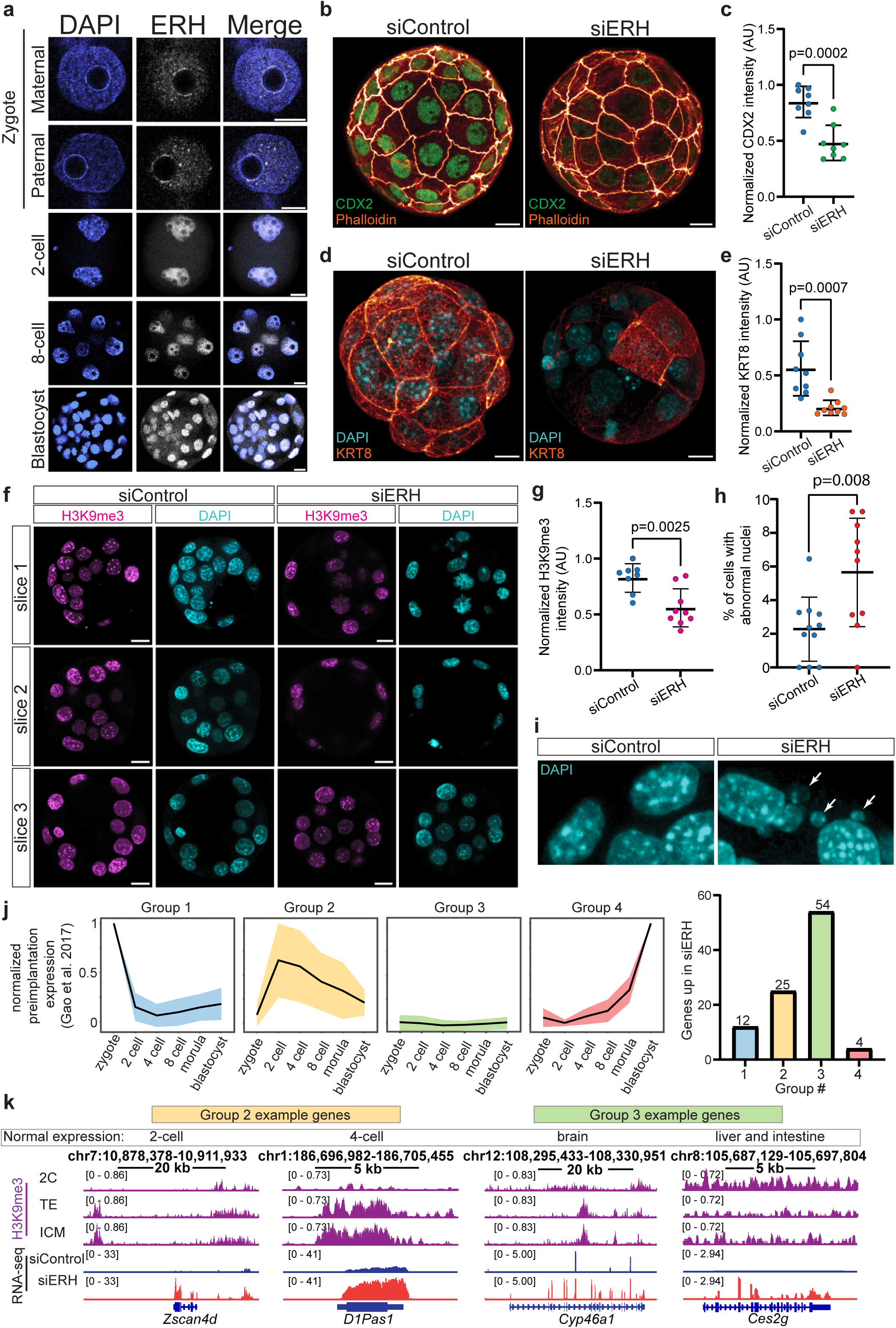
ERH is expressed during pre-implantation mouse development and is critical for normal blastocyst development by repressing 2-cell and alternative lineage genes. **a.** ERH (white) and DAPI (blue) staining during mouse pre-implantation development. Representative confocal z-sections are shown (zygote n=15, N=3, 2-cell stage n=18, N=3, 8-cell stage n=6, N=2, blastocyst n=10, N=3). Scale bars, 10µm. **b.** Representative images of trophectoderm marker Cdx2 (green) in siControl and siERH blastocysts with Phalloidin (orange) marking cell boundaries. Scale bars, 10µm. **c.** Quantification of CDX2 intensity in siControl vs siERH blastocysts (unpaired t-test, n=8 blastocysts for each siRNA). **d.** Representative image of trophectoderm marker KRT8 (orange) in siControl and siERH blastocysts with DAPI (blue). Scale bars, 10µm. **e.** Quantification of KRT8 intensity in siControl vs siERH blastocysts (unpaired t-test, n=9 blastocysts, n=8 blastocysts for each siRNA). **f.** Representative images of H3K9me3 (magenta), DAPI (teal) in siControl vs siERH blastocysts. Scale bars, 10µm. **g.** Quantification of H3K9me3 level in siControl vs siERH blastocysts. (unpaired t-test, n=8 for siControl and n=9 for siERH blastocysts). **h.** Percentage of cells per blastocyst with abnormal nuclei. (unpaired t-tes, n=11 for siControl and n=9 for siERH blastocysts). **i.** Representative image of DAPI staining siCTRL and siERH blastocysts with white arrows indicating DNA shedding. **j.** ERH repressed blastocyst genes (adjusted p-value<=0.05, log2fc>=0.5) grouped based upon normal expression patterns exhibited during mouse pre-implantation development re-analyzed from Gao et al., 2017^33^. **k.** Example gene tracks of genes significantly upregulated in siERH vs siControl blastocysts (adjusted p-value<=0.05, log2fc>=0.5, n=3) with H3K9me3 tracks for 2 cell, trophectoderm and ICM cells from^11^. See also Extended Data 5 and Supplementary Tables 5,6.

To investigate if ERH is involved in pre-implantation lineage specification, we depleted ERH levels by siRNA micro-injection at the 1-cell stage of mouse embryos, compared to a nontargeting siRNA control, and assessed impacts on the blastocysts (Extended Data Fig. 5c). ERH depletion did not cause major morphological changes in blastocyst structure, as both siControl and siERH blastocysts form a cavity and segregate ICM and trophectoderm cells (Extended Data Fig. 5c,d). Staining blastocysts for key transcription factors involved in lineage specification revealed similar levels of the pluripotency marker OCT4 in ICM and TE cells between siControl and siERH blastocysts (Extended Data Fig. 5e). However, the levels of the trophectoderm-determining transcription factor CDX2^34^ (Fig. 3b,c) and the trophectoderm marker Keratin 8^35^ (Fig. 3d,e) were significantly decreased in the ERH depleted blastocysts. Thus, ERH is required for proper trophectoderm differentiation.

The role of ERH in H3K9me heterochromatin regulation in *S. pombe*^1,16^ and human fibroblasts^2^ suggests that defects in trophectoderm differentiation may arise from alterations in gene silencing mechanisms. Indeed, we observed a 32% decrease (p=0.0025) in H3K9me3 in blastocysts of ERH-depleted embryos (Fig. 3f,g and Supplementary Table 5). We also observed a significant increase in nuclear DNA shedding in the siERH blastocytes (Fig. 3h,i; Extended Data Fig. 5f), similar to the observed nuclear DNA shedding in siERH hESCs (Extended Data Fig. 1g). Decreased keratin levels as observed in siERH blastocytes (Fig. 3d,e) have been shown to make trophectoderm cells of blastocyts more susceptible to DNA shedding^36^, as has decreased H3K9me3^18^.

To characterize how these changes impact gene expression, we performed RNA-seq on mouse blastocysts 4 days after injecting siControl or siERH at the 1 cell stage (Supplementary Table 6), which confirmed an ∼80% knockdown of ERH (Extended Data Fig. 5g). We classified genes upregulated in ERH depleted blastocysts into 4 groups, based upon their expression during normal mouse preimplantation development^33^ (Fig. 3j; Supplementary Table 7). Group 1 consists of transcripts present at the zygote stage, including maternally inherited transcripts, that become silenced or degraded as development progresses. Group 2 consists of transcripts that are highly expressed in the 2 to 8 cell stages and low or absent in the zygote and blastocyst stage. Group 3 consists of transcripts that are absent or lowly expressed at all stages measured. Group 4 consists of transcripts with the highest expression in the blastocyst stage. Of the ERH-repressed genes (i.e., genes whose expression goes up upon ERH knockdown), 25 normally have peak expression during the 2 to 8 cell stage (group 2) and 54 are normally not expressed during preimplantation development (group 3). ERH-repressed group 2 genes included the 2-cell specific *Zscan4d*^37^ and the 4-cell specific *D1Pas1*^38^ (Fig. 3k). ERH-repressed group 3 alternative lineage genes included genes specific to the brain^39^, liver, and intestine ^40^ (Fig. 3k).

Comparing our RNA-seq data with existing mouse preimplantation H3K9me3 ChIP-seq data^11^, we found that the genes repressed by ERH at the 2-cell stage and later, alternative lineage genes were initially marked by H3K9me3 (Fig. 3k). We further found that upon ERH depletion, key markers of primitive endoderm, including *Gata4* and *Sox17*^41^, were significantly repressed (Extended Data Fig. 5h), while epiblast genes were unchanged (Extended Data Fig. 5i). We conclude that ERH is necessary to silence 2-8 cell stage genes and prevent the spontaneous activation of later developmental genes during the blastocyst stage, and that failure to silence results in improper lineage specification in the blastocyst.

### ERH is necessary for proper lineage commitment and restriction during human ESC differentiation

Our finding that ERH is critical for blastocyst development led us to ask if ERH depletion impacted the ability of hESCs to differentiate. When we used Activin A and the GSK-3 Inhibitor CHIR99021 together to differentiate hESCs to an endoderm identity^42^, as expected within 24 hours we observed repression of pluripotency factors and initial expression of endodermal markers in a subset of cells (Extended Data Fig. 6a). Yet depletion of ERH in H1 hESCs followed by 24 hour endoderm differentiation showed a failure to decrease NANOG protein levels (Fig. 4a and Extended Data Fig. 6b) and a reduced emergence of SOX17+ cells (Fig. 4a). To assess if this observation was specific to endoderm, we also performed differentiation to ectoderm, with control and ERH depleted cells, for 2 days with retinoic acid treatment. We observed sustained expression of the pluripotency factor OCT4 and decreased abundance of the ectodermal transcription factor PAX6 (Fig. 4b).

**Figure 4.**
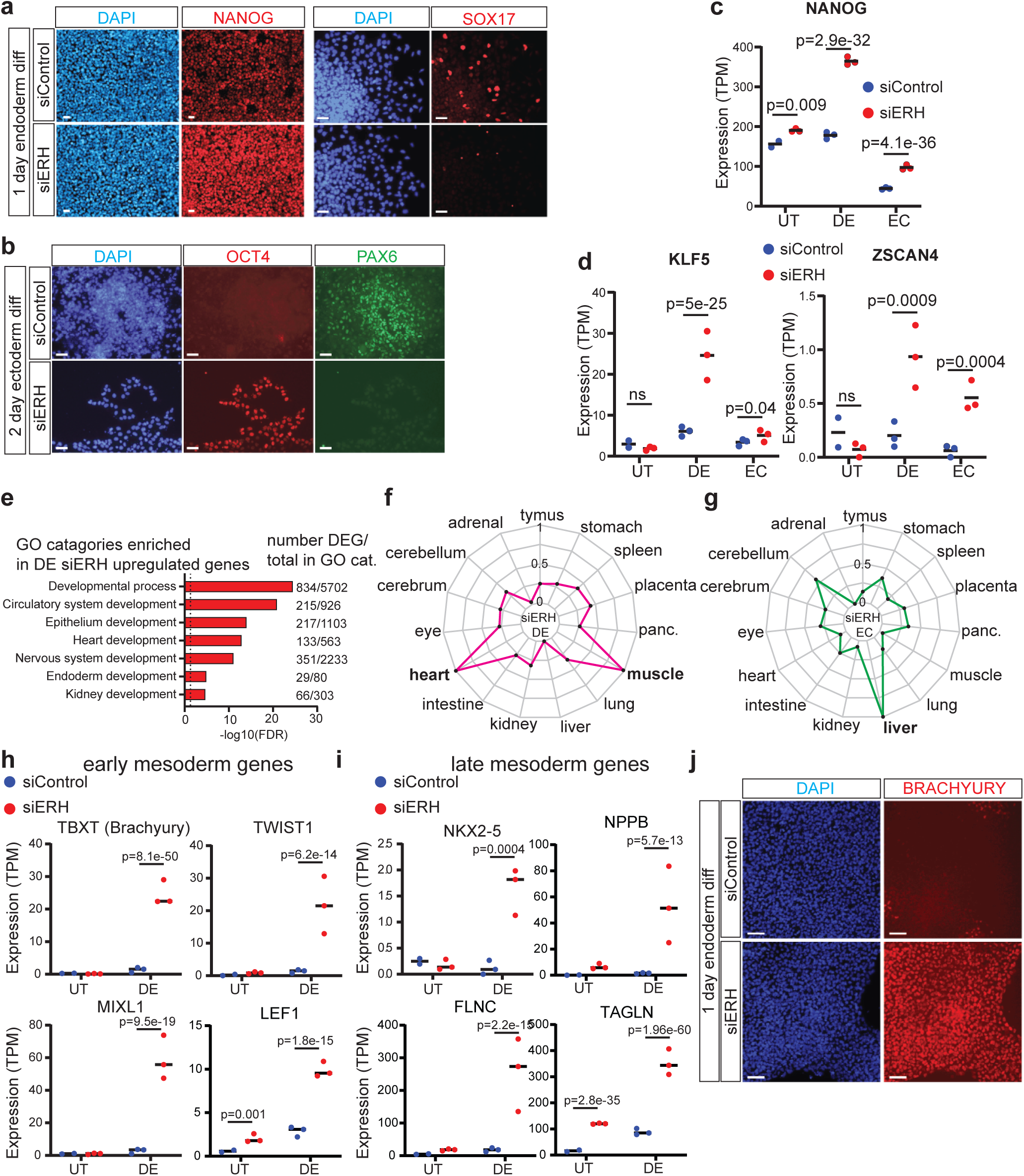
ERH represses pluripotency and alternative lineage gene activation during hESC differentiation. **a.** Representative images of NANOG (red), SOX17 (red) and DAPI (blue) during 1 day of endoderm differentiation of siControl and siERH H1 hESCs. Scale bars, 50µm. **b.** Representative images of OCT4 (red), PAX6 (green) and DAPI (blue) during 2 days of ectoderm differentiation of siControl and siERH H1 hESCs. Scale bars, 50µm. **c.** Plot of NANOG expression in transcripts per million (TPM) by RNA-seq in siControl and siERH treated H1 hESCs cultured in pluripotency (H1), 1 day endoderm differentiation (DE) and 2 day ectoderm differentiation (EC). (DESeq2 adjusted p-value reported). **d.** Plot of KLF5 and ZSCAN4 expression in transcripts per million (TPM) by RNA-seq in siControl and siERH treated H1 hESCs cultured in pluripotency (H1), 1 day endoderm differentiation (DE) and 2 day ectoderm differentiation (EC). (DESeq2 adjusted p-value reported). **e.** GO analysis (PantherDB, statistical overrepresentation) of genes upregulated (padj<=0.05, log2 fold change >= 0.5) in siERH vs siControl H1 hESCs 1 day endoderm differentiation. **f.** Radar plot showing cumulative expression profile across indicated human tissues, data downloaded from BBI Descartes database^77^, for genes upregulated (padj<=0.05, log2 fold change >= 0.5) in siERH vs siControl H1 hESCs 1 day endoderm differentiation. **g.** Radar plot showing cumulative expression profile across indicated human tissues, data downloaded from BBI Descartes database^77^, for genes upregulated (padj<=0.05, log2 fold change >= 0.5) in siERH vs siControl H1 hESCs 2 day ectoderm differentiation. **h.** Example early mesodermal genes activated during endoderm differentiation by ERH depletion, transcripts per million (TPM) values from RNA-seq replicates shown (DESeq2 adjusted p-value). **i.** Example late mesodermal genes activated during endoderm differentiation by ERH depletion, transcripts per million (TPM) values from RNA-seq replicates shown DESeq2 adjusted p-value). **j.** Representative image of DAPI (blue), BRACHYURY (red) of siControl and siERH treated H1 hESCs after 1 day of endoderm differentiation. Scale bars, 50µm. For all RNA-seq TPM values n=3, except for siCTRL UT where n=2. See also Extended Data Fig. 6 and Supplementary Tables 7,8.

To investigate transcriptional changes caused by ERH depletion, we performed RNA-seq of siRNA control and ERH depleted H1 hESCs (Extended Data Fig. 6c and Supplementary Table 8) during pluripotent culture, after 1 day of endoderm or 2 days of ectoderm differentiation. While ERH depletion in fibroblasts+OSKM resulted in more gene activation than repression (Fig. 2c; Extended Table 3), depletion in pluripotent hESCs exhibited less derepression (Extended Data Fig. 4d; Extended Table 8), consistent with the difference in the importance of ERH in H3K9me3 maintenance between fibroblasts and hESC state (Fig. 1). Genes normally repressed in the siControl endoderm and ectoderm differentiation were preferentially upregulated by ERH depletion (endoderm, Extended Data Fig. 6f; ectoderm Extended Data Fig. 6g).

During endoderm differentiation, we observed that ERH depletion preferentially upregulated genes identified as higher in naïve vs primed hESCs^43^ (Extended Data Fig. 6h), beyond what was observed in the pluripotent state (Extended Data Fig. 6e). For example, the gene for the pluripotency factor NANOG, previously demonstrated to be repressed by H3K9me3 during pluripotency exit^44^, was significantly upregulated by ERH depletion (Fig. 4c), consistent with our immunostaining (Fig. 4a and Extended Data Fig. 6b). Additionally, genes found to be expressed in human pluripotent stem cells were upregulated during endoderm and ectoderm differentiation after ERH depletion, including *DUSP5/6, NODAL, LEFTY1/2*, and *DACT1* (Extended Data Fig. 6i and Supplementary Table 8). Genes associated with the 2 cell stage and naïve pluripotency, including *ZSCAN4*^37^*, KLF5*^31^*, ESRRB*^45^*, NR5A2*^46^ and *NANOGP1*^47^, were upregulated during siERH-treated hESC differentiation (Fig. 4d and Extended Data Fig. 6j). The expression of naïve genes after ERH depletion was at a much lower level than in true naïve hESCs (Extended Data Fig. 6k), indicating permissive expression but not spontaneous primed to naïve conversion. We conclude that ERH is required to repress pluripotency genes during hESC pluripotency exit.

During differentiation, a key function of H3K9me3 heterochromatin is to repress alternative lineage genes^3,11^. Our finding that ERH depletion in blastocysts upregulates lineage-incompatible genes (Fig. 3j,i) led us to investigate if a similar consequence occurs during hESC differentiation. GO analysis of genes upregulated by ERH depletion during endoderm and ectoderm differentiation confirmed activation of transcriptional networks typically not induced by the specific differentiation treatment (Fig. 4e, Extended Data Fig. 7a and Supplementary Table 10). ERH depletion upregulated meiotic and gametogenic genes in all tested developmental stages (Extended Data Fig. 7b), consistent with previous findings in human fibroblasts^2^ and the role of ERH in S. pombe sporulation^1^. To further assess the cell lineages in which the upregulated genes function, we generated composite expression profiles of the genes upregulated by ERH depletion during endoderm (Fig. 4f) and ectoderm differentiation (Fig. 4g) from expression data downloaded from the BBI Descartes human single cell atlas^48^, and found that heart, muscle and liver genes are preferentially activated. Directed differentiation of hESCs to endoderm^42^, instead induced both early and late mesodermal genes^49^ when ERH was depleted (Fig. 4h,i). Notable inappropriate gene activation in siERH definitive endoderm included *NXK2-5, NPPB* and *FLNC*, typically expressed in heart muscle, as well as *TAGLN,* a canonical smooth muscle marker (Fig. 4i). Increased levels of BRACHYURY protein, a key determinant of early mesoderm^50^, was also confirmed by immunofluorescence (Fig. 4j). Inappropriate activation during ectoderm differentiation with ERH knockdown included genes typically expressed in liver development (Extended Data Fig. 7c). Thus, while ERH appears dispensable to maintain hESC pluripotency, it is essential to restrict cell identity for proper germ layer differentiation.

### ERH represses transposable elements and proximal genes

Having observed that ERH represses pluripotency and alternative lineages during hESC differentiation, we proceeded to investigate how this repression was elicited. During pluripotency-to-endoderm differentiation at protein-coding genes that are upregulated in siERH DE vs siControl DE, H3K9me3 levels only slightly increase in and around those genes (i.e., without siERH) during endoderm differentiation (Extended Data Fig. 8a). However, repeat elements including L1 LINEs do normally exhibit a substantial H3K9me3 gain during endoderm differentiation (Fig. 5a). Repression of transposable elements is a canonical function of H3K9me3 heterochromatin^51^ and transposable elements can regulate developmental gene networks by acting as enhancers for transcription factor binding^31,52–55^. We therefore assessed transposable elements in our RNA-seq data from siControl and siERH-treated H1 hESCs during self-renewal and differentiation (Supplementary Table 9), with the hypothesis that dysregulation of specific transposable elements might underlie the observed differentiation defects in siERH treated hESCs.

**Figure 5.**
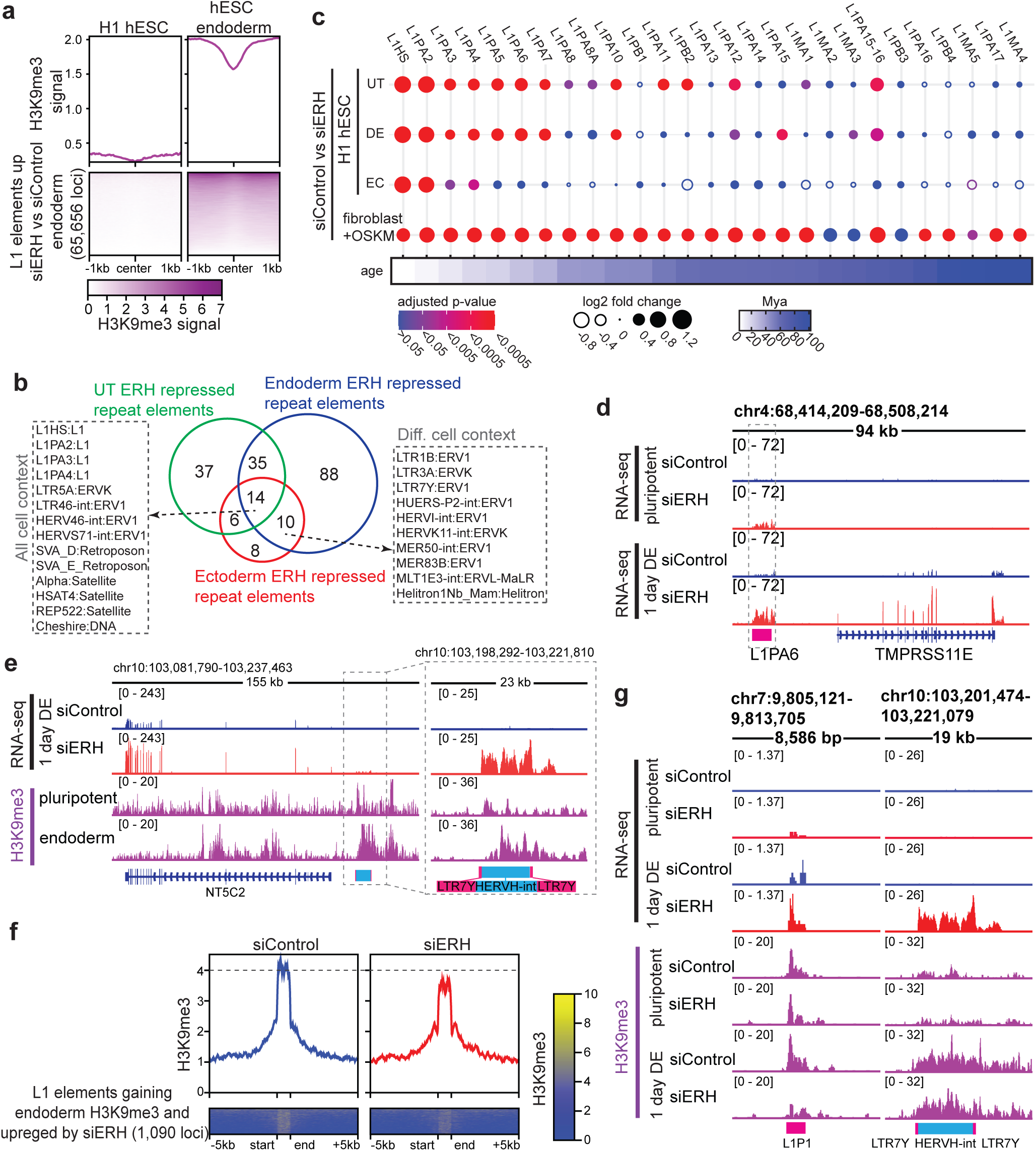
ERH represses evolutionarily young transposable elements with regulatory function. **a.** H3K9me3 data from H1 hESCs (GSM1003585) and hESC derived endoderm (GSM916057) at 65,656 L1 element loci +/- 1kb upregulated (adjusted pvalue<=0.05; log2foldchange>=0.5) by ERH depletion during 1 day endoderm differentiation. **b.** Venn diagram showing intersect of repeat classes upregulated (TETranscripts, padj<0.05, log2fc>0.5) by ERH depletion in H1 hESCs untreated (UT), 1 day endoderm (DE) or 2 day ectoderm (EC) conditions. **c.** Expression change and significance, TEtranscripts^78^ analysis, of L1 LTR transposable elements in ERH depletion in H1 hESCs untreated (UT), 1 day endoderm (DE) or 2 day ectoderm (EC) listed in order of evolutionary age as determined in^59^. **d.** Example browser track showing expression of L1 element and proximal gene in siControl and siERH treated H1 hESCs in the pluripotent and 1 day DE conditions. **e.** Example browser track of gene with promoter proximal repeat element gaining H3K9me3 during endoderm differentiation that is upregulated by ERH depletion. **f.** H3K9me3 CUT&RUN signal at L1 elements that exhibited 2-fold RNA increase in siERH vs siControl H1 hESC endoderm and switched from H3K9me3 unmarked in pluripotent to marked in endoderm. **g.** Browser track showing RNA-seq of example repeat element and nearby genes in siControl and siERH treated H1 hESCs after 1 day of endoderm differentiation and H3K9me3 tracks from H1 hESCs (GSM1003585) and hESC derived endoderm (GSM916057). See also Extended Data Fig. 7,8 and Supplementary Table 10.

We found that transposable elements were upregulated by ERH knockdown in H1 hESCs, definitive endoderm, and ectoderm, including classes of L1 elements and the hominid-specific^56^ SINE VNTR and Alu (SVA) elements (Fig. 5b; Extended Data Fig. 8b). Satellite repeats were also derepressed (Fig. 5b), as were morula-blastocyst specific *LTR7Ys*^31^ and human 8 cell stage expressed *HERVKs*^57^ (Fig. 5b; Extended Data Fig. 8c,d), which act as transcription factor binding sites for pluripotency^52,58^ and lineage specific genes^55^. After stratifying L1 elements by their evolutionary age^59^, we found that ERH preferentially represses evolutionarily young L1 repeats in pluripotent and differentiating hESCs, but broadly represses L1 elements in terminally differentiated fibroblasts (Fig. 5c). By filtering our data for genes expressed during pluripotency that exhibit ERH-dependent repression during differentiation, we uncovered evolutionarily young transposable elements (Fig. 5d and Extended Data Fig. 8e,f) and L1 repeats within 25 kb (Extended Data Fig. 8g) in regions normally marked by H3K9me3 in hESC-derived endoderm (Fig. 5e note different scales on subset to right; Extended Data Fig. 9a).

To assess if repeat expression upon ERH knockdown during endoderm differentiation was due to H3K9me3 loss, we performed H3K9me3 CUT&RUN in siControl and siERH hESCs differentiated to endoderm. We found the increased L1 element RNA expression upon ERH depletion (Fig. 5c) did not globally correspond to a loss of H3K9me3 (Extended Data Fig. 9b). However, when we restricted the analysis to L1 TEs that gained H3K9me3 with at least a 2-fold increase in RNA in siERH endoderm, the L1 loci showed decreased H3K9me3 upon ERH depletion (Fig. 5f). We found repeat element loci with increased RNA expression with (Fig. 5g left) and without H3K9me3 loss (Fig. 5g right) indicating that RNA expression from TEs upon ERH depletion during endoderm differentiation was not necessarily dependent upon H3K9me3 loss. We observe instances of endoderm specific upregulation of genes proximal to evolutionarily young repeats, such as L1 and LTR7Ys, with reduced H3K9me3 in ERH depleted endoderm (Extended Data Fig. 9c). ERH therefore represses evolutionarily young and lineage-specific repeats, thus preventing improper activation of pluripotency and alternative lineage genes during differentiation to maintain lineage specification integrity. The role of H3K9me3 and ERH at this cell fate change likely occurs predominantly at transposable element regions and not gene bodies.

### ERH is recruited to chromatin by cell-specific cofactors

Our observation that ERH is required for maintenance of H3K9me3 in fibroblasts (Fig. 1b) but dispensable for hESC H3K9me3 (Fig. 4b) led us to investigate potential ERH protein interactions that may drive the difference in regulatory mechanism. We focus the experiments on the pluripotent and terminally differentiated states as they exhibit contrasting requirements for ERH in H3K9me3 maintenance (Fig. 1b-g) and very different H3K9me3 distribution patterns (Fig. 1d, i; Extended Data Fig 1g). In *S. pombe* Erh is recruited by the RNA binding factor Mmi1^1^. Mmi1 itself is not seen in mammalian cells^60,61^, but since it is a YTH domain-containing RNA binding factor, we investigated potential interactions of ERH with human YTH domain proteins YTHDC1 and YTHDC2. YTHDC1 regulates H3K9me3 through SETDB1 recruitment and repression of repeat elements in mouse ESCs^62,63^ and YTHDC2 has been shown to positively regulate transposable elements and the non-coding RNA LINC-ROR in human hESCs^64^. To specifically interrogate on chromatin ERH interacting proteins, we isolated soluble nucleoplasm (NPL) and insoluble chromatin-associated fractions (CAF) and then performed immunoprecipitation of ERH (Fig. 6a). In fibroblasts but not hESCs, we observed that ERH interacts with YTHDC1 in the chromatin fraction (Fig. 6b,c). ERH did not interact with YTHDC2 in the chromatin fraction in either hESCs or fibroblasts, despite being present in both cell types (Fig. 6b,c). We also probed for SAFB1, a known ERH interacting protein^65^, and observed consistent ERH-SAFB1 interactions in both fibroblasts and pluripotent hESCs (Fig. 6b,c), demonstrating that YTHDC1 but not SAFB1 interactions with ERH are cell type-dependent. To investigate the cause of this differential interaction we first analyzed ERH levels, finding that while ERH RNA is not significantly different between pluripotency and fibroblasts (Extended Data Fig. 10a) or between pluripotent and differentiated hESCs (Extended Data Fig. 6c), fibroblasts stain more strongly for ERH protein (Extended Data Fig. 10b). We observe that ERH protein levels are higher in cells on the edge of hESC colonies (Extended Data Fig. 10c) and become higher in all cells after 1 day of endoderm differentiation (Extended Data Fig. 10c). These results indicate that ERH involvement in H3K9me3 regulation and RNA binding protein regulation may be regulated by ERH protein level.

**Figure 6.**
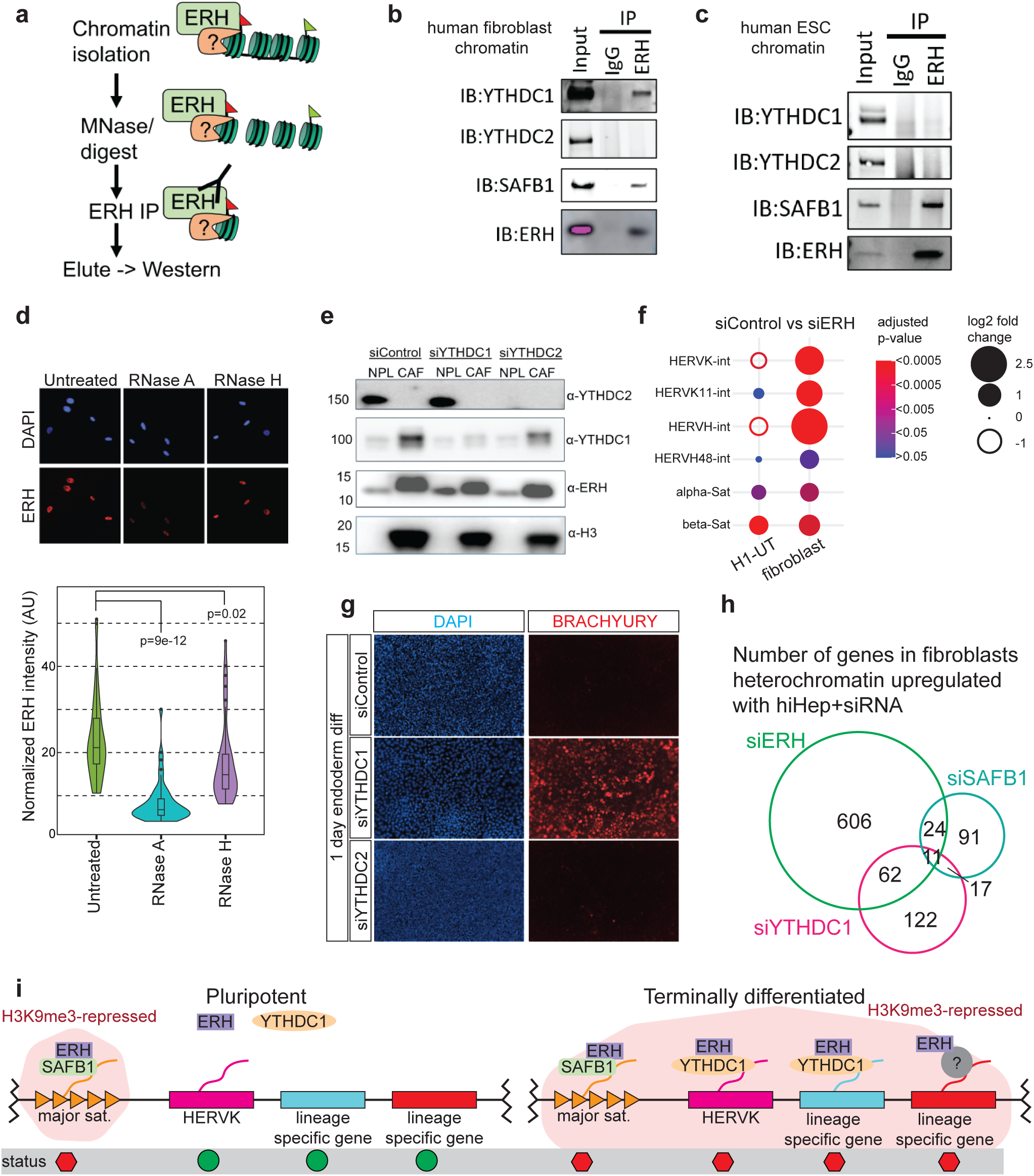
Cell type specific and independent recruitment of ERH to chromatin by RNA binding proteins regulates gene and repeat repression. **a.** Strategy for identifying chromatin-bound ERH interaction partners by isolation of soluble nucleoplasm and chromatin-associated protein fractions prior to immunoprecipitation. **b.** Immunoprecipitation (IP) of ERH from the chromatin fraction of human fibroblasts. Western blot analysis of YTHDC1, YTHDC2, SAFB1, and ERH in the input and IP samples. **c.** Immunoprecipitation (IP) of ERH from the chromatin fraction of human ESCs. Western blot analysis of YTHDC1, YTHDC2, SAFB1, and ERH in the input and IP samples. **d.** Immunofluorescence (IF) of ERH after RNase A and RNase H treatment. **e.** Western blot of ERH, YTHDC1, and YTHDC2 in nucleoplasm and chromatin-associated protein fractions of human fibroblasts after siYTHDC1 and siYTHDC2 treatment. **f.** Expression change and significance, TEtranscripts^78^ analysis, of HERVK HERVH and satellite repeats in hESCs and human fibroblasts after siERH. **g.** Immunofluorescence (IF) of BRACHYURY in siCTRL, siYTHDC1 and siYTHDC2 after 1 day endoderm differentiation. **h.** Venn diagram of gene coregulation after siERH, siYTHDC1 and siSAFB1 in human fibroblasts after addition of hepatic reprogramming factors, FOXA3, HNF1A, HNF4A. **i.** Proposed model for cell-type specific, and cell-type agnostic gene and repeat regulation by ERH, YTHDC1, and SAFB1. See also Extended Data Fig. 9

As S. pombe Erh1 chromatin recruitment is RNA and Mmi1 dependent, we next tested if chromatin association of ERH was RNA dependent and if either YTHDC1 or YTHDC2 facilitated the recruitment. We employed an immunostaining approach^66^ in which permeabilized nuclei were treated with RNase A and then stained for ERH, and found that ERH nuclear retention was significantly decreased (Fig. 6d). In contrast, RNase H treatment, which digests DNA:RNA hybrids, did not release ERH from the nucleus (Fig. 6d). In an orthogonal approach, RNase A treatment of fibroblasts during soluble nuclear protein isolation released a portion of chromatin-bound ERH into the nucleoplasm, indicating that its chromatin retention is at least partially RNA-dependent (Extended Data Fig. 10d). To determine if YTHDC1 or YTHDC2 facilitate ERH chromatin recruitment, we targeted each separately with siRNAs in human fibroblasts and observed that YTHDC1 but not YTHDC2 depletion led to decreased levels of ERH in the chromatin fraction and increased ERH in the soluble nuclear fraction (Fig. 7e). We confirmed the role of YTHDC1 in ERH retention utilizing our cell permeabilization and immunostaining assay (Extended Data Fig. 10e). From these experiments we concluded that ERH associates with chromatin with YTHDC1 in fibroblasts, but not in hESCs, and that a subset of ERH protein requires RNA and YTHDC1 for chromatin localization.

We next assessed for co-regulation of common targets by ERH, YTHDC1, and YTHDC2, first by evaluating expression changes of published YTHDC1 and YTHDC2 targets, ERVK elements repressed by YTHDC1^67^, and HERVH elements activated by YTHDC2^64^. ERH represses HERVK and HERVH elements in fibroblasts treated with OSKM factors but not in pluripotent hESCs (Fig. 6f), matching the interaction pattern of ERH with YTHDC1 (Fig. 6b,c). Major satellite repeats, which are regulated by SAFB1^68^, are repressed by ERH in both fibroblasts and pluripotent hESCs (Fig. 6f).

As we found that ERH interacts with YTHDC1 in differentiated cells but not pluripotent cells, we investigated if YTHDC1 depletion recapitulates aspects of ERH depletion during hESC differentiation. Depletion of YTHDC1 but not YTHDC2 during endoderm differentiation of hESCs resulted in an increase of Brachyury staining indicative of a mesodermal fate (Fig. 6g), matching our findings from ERH depletion during endoderm differentiation (Fig. 5f). We found that ERH represses many genes not regulated by its interacting partners YTHDC1 and SAFB1, which predominately repress different subsets of ERH targets (Fig. 6h). Taken together, the results indicate that various RNA binding proteins function non-redundantly to recruit ERH to different targets. Based upon these findings, we propose that ERH interactions with RNA binding proteins, including YTHDC1 and SAFB1, facilitate cell-specific repression of genes and repeat elements (Fig. 6i).

## Discussion

We discovered that depletion of ERH reverts the H3K9me3 landscape in terminally differentiated cells to an hESC-like state, permissive to activation of early embryonic genes. This surprising result led us to investigate the role of ERH in establishing gene repression during the earliest stages of mammalian lineage segregation, in maintaining repression of pluripotency, and enabling germ layer specification. We found that ERH controls H3K9me3 for repression of naïve and alternative lineage genes during the cell fate commitment of totipotent cells to ICM and trophectoderm in mouse blastocysts as well as of hESCs to the endoderm and ectoderm cell lineages, to enforce cell fate and prevent anomalous gene activation. Between the three systems we employed, mouse preimplantation development, hESC differentiation, and fibroblast iPSC reprogramming, we observe that ERH commonly repressed a shared set of naïve genes such as *ZSCAN4* across all systems, indicating that ERH is involved in both the initial repression of this early transcriptional program and in maintaining repression in terminally differentiated cells. Remarkably, ERH is the sole apparent mechanism for repressing meiotic genes in *S. pombe,* is conserved in repressing germ cell genes in mammals^2^, and has expanded to foundationally control diverse other types of developmental genes and stages.

In a developmental context, the failure to repress 2-cell, 8-cell and pluripotency genes after ERH knockdown results in delays of specific-lineage gene expression, including primitive endoderm genes in mouse blastocysts and various endoderm and ectoderm genes during hESC differentiation. Furthermore, we demonstrated that ERH is necessary to maintain repression of alternative lineage genes, as ERH depletion elicited precocious expression of genes typical of later cell differentiation in mouse blastocysts and human hESCs, as well as heterochromatinized genes during initial iPSC reprogramming. We observed that evolutionarily young transposable elements, including the human-specific *LTR7Ys*^31^, are especially upregulated by ERH depletion during hESC differentiation and iPSC reprogramming. Much of the lineage specific expression changes upon ERH depletion during hESC differentiation may be attributed to increased accessibility of transcription factor binding sites among derepressed repeats^31,52,55^.

A key finding is that ERH depletion in human fibroblasts reverted the H3K9me3 chromatin landscape to a hESC-like state. The reversion to a hESC-like H3K9me3 landscape permits binding of SOX2 in fibroblasts to endogenous hESC SOX2 targets, activating embryonic genes. That ERH is necessary for maintaining the broad H3K9me3 domains observed in somatic cells, but is dispensable for maintaining hESC H3K9me3 peaks, is consistent with our observation that ERH depletion in hESCs did not change global H3K9me3 levels. We suggest that the residual H3K9me3 peaks in pluripotent cells, enriched for zinc finger motifs and KAP1 binding, may be H3K9me3 nucleation sites and that, during differentiation, ERH is essential for the spread of H3K9me3 from these sites. Importantly, when ERH is knocked down in fibroblasts, we observe increased activation of many genes and transposable elements that harbor a residual H3K9me3 peak, yet lose broad H3K9me3, indicating that residual H3K9me3 peaks are not sufficient to block gene activation. Our knowledge of H3K9 methylation nucleation and spread has been primarily elucidated through studies in *S. pombe*^69–71^, leaving further investigation needed into how H3K9me3 nucleation and spread occurs in multicellular organisms.

We identify that ERH interacts with and is partially recruited to the chromatin by the RNA binding protein YTHDC1 in somatic cells, but not in pluripotent hESCs. We find that ERH and YTHDC1 repress similar targets in fibroblasts and differentiating hESCs, but not in homeostatic pluripotent hESCs. In contrast, we observe that ERH interacts with the RNA binding protein SAFB1 in both hESCs and human fibroblasts and likewise both repress the satellite repeat expression in hESCs and fibroblasts. Our knockdowns of YTHDC1 and gene expression analysis of YTHDC1 and SAFB1 fail to fully explain how ERH is recruited to chromatin to repress transcription. We thus speculate that ERH is recruited to cell-type specific genes and repeats through a diverse network of chromatin-associated RNA binding factors that are yet to be fully identified.

We observe in mouse development and hESC systems that ERH is functionally required in multiple developmental contexts to repress meiotic and gametogenic genes, as it is in *S. pombe*^1^, although the gene members targeted by ERH has greatly evolved and expanded in mammals. Our findings on meiotic gene repression are supported by the recent identification of ERH in a screen for proteins silencing germline genes in mESCs^72^. Other aspects of *Erh1* function in *S. pombe,* such as the targeting of RNAs for degradation by the RNA exosome^1^, appear to be conserved as we observed upregulation of developmental genes typically downregulated by RNA exosome degradation^73^. ERH was recently found by mass spectrometry to associate with the Microprocessor and NEXT complexes in mouse ESCs^74^. Evolutionarily divergent functions of ERH may be driven by evolutionarily young transposable elements that retain RNA expression, similar to how transposable elements have influenced the evolution of KRAB Zinc finger proteins DNA dependent targeting of H3K9me3^75,76^.

We found that ERH enacts gene repression at the earliest stages of development and that it is essential for maintaining this repression in terminally differentiated cells. Our findings advance the understanding of how heterochromatin-based repression is established and maintained during development. We propose that understanding mechanisms by which ERH is employed to direct H3K9me3 to specific loci in a developmentally coordinated fashion will be crucial to modulating developmental H3K9me3 rearrangements to achieve greater control of cell identity.

## Methods

### Cell lines culture and condition

Human H1 ES cells were obtained from WiCell (WA01), grown on Matrigel coated wells and cultured in mTeSR1 (Stem Cell Tech, 85850) medium at 37°C and 5% CO_2_. Medium was changed daily. Human BJ foreskin fibroblasts were obtained from ATCC (CRL-2522) and cultured in Eagle’s Minimum Essential Medium (EMEM, Sigma-Aldrich M2279) supplemented with 10% fetal bovine serum (FBS, Hyclone SH30071) and 2mM Glutamax (ThermoFisher 35050061) at 37°C and 5% CO_2_.

### Mouse embryo work

Mouse embryo experimentation was approved by the Biological Resource Center Institutional Animal Care and Use Committee (IACUC), Agency for Science, Technology and Research (IACUC Protocol 806983). C57BL/6 wild-type 3–4-week-old female mice were superovulated using 5 IU of pregnant mare serum (PMS, National Hormone and Peptide Program) gonadotropin given intraperitoneally and 5 IU of recombinant chorionic gonadotrophin (CG, National Hormone and Peptide Program) given 48 h after and immediately before mating, according to animal ethics guidelines of the University of Pennsylvania. Embryos were flushed from oviducts of plugged females using M2 medium (Merck) and cultured in KSOM+AA (Merck) covered by mineral oil (Sigma, M-5310), at 37 °C and 5% CO2.

### Mouse embryo immunofluorescence and microscopy

Embryos were fixed in 4% paraformaldehyde (Electron Microscopy Sciences, 15710) for 30 min at room temperature washed twice in PBS with 0.1% Triton X-100 (Millipore Sigma, T8787), permeabilized for 20 min in PBS with 0.5% Triton X-100, and blocked in 3% bovine serum albumin (Gemini Bio, 700-100P) for 30 min. Embryos were then incubated at 4 °C overnight in primary antibodies diluted in blocking solution at the following concentrations: ERH (Abcam, ab166620) at 1:200, Keratin-8 (DSHB, THROMA-I) at 1:20, H3K9me3 (Abcam, ab8898) at 1:1000, and CDX2 (Invitrogen, ZC007) at 1:200. After primary antibody incubation, embryos were washed 5 times for 20 min in PBS with 0.1% Triton X-100 and incubated for 30 min in Alexa Fluor-conjugated secondary antibodies (Invitrogen, A32731, A32723, A32733, A32728 and A11006) diluted in blocking solution to 1:500. Phalloidin-Alexa Fluor 555 (Invitrogen, A34055) diluted to 1:500 and DAPI (Sigma, 10236276001) diluted to 1:500 in blocking solution were also used to label F-actin and DNA, respectively. Embryos were washed three times in PBS with 0.1% Triton X-100 before mounting in PBS covered with mineral oil in an 8-well μ-Slide (Ibidi, #80827). Embryos were imaged using a laser-scanning confocal microscope (Leica SP8) with a water-immersion Apochromat 40× 1.1 NA objective and HyD detectors. Samples were excited with standard laser lines (405-nm, 488-nm, 552-nm, 638-nm).

### Mouse embryo siRNA injection experiments

Microinjections were performed using a FemtoJet (Eppendorf). For ERH knockdown experiments, mouse embryos were injected with two siRNAs (Qiagen, S104403294 & S104403287) against ERH at a total concentration of 400 nM. For knockdown experiments AllStars negative control siRNA (Qiagen, 1027281) was injected in embryos at a concentration of 400nM.

### Differentiation of hESCs

Prior to differentiation, for samples to be analyzed by RNA-seq, cells grown to a density of 80% were digested with Accutase (Stem Cell Technologies, 07920) for 7 min. Then, DMEM/F12 was added and gently pipetted to generate a cell suspension. After centrifugation, cells were resuspended in mTeSR1 medium with 5uM ROCK inhibitor Y-27632 (Cayman Chemical, 10005583) and seeded at 8X10^5^-1X10^6^ cells per well in a 6 well plate coated with Matrigel (Millipore Sigma, CLS354277) 24 hours prior to differentiation treatment. For definitive endoderm differentiation of hESCs, mTeSR1 medium was removed and cells were cultured in DMEM/F12 (Thermo Fisher, 11320033) supplemented with 10% (v/v) Fetal Bovine Serum (Hyclone, SH30071) with 100ng/ml Activin A (Peprotech, 120-14P) and 3uM CHIR99021 (Cayman Chemical, 13122). For ectoderm differentiation of hESCs, mTeSR medium was removed and cells were cultured in DMEM/F12 with 5uM Retinoic Acid (Cayman Chemical, 11017).

### Cell culture siRNA transfection experiments

All ERH knockdown experiments were performed with two cycles of siRNA transfection (Silencer Select siRNA, Thermo Fisher, s4815), three days apart. Transfections were performed in fibroblasts using 5nM final concentration of siRNA and in hESCs using a 10nM final concentration of siRNA and 2.5ul/ml final concentration of Lipofectamine RNAiMAX (Thermo Fisher #31985070).

### Reprogramming factor expression experiments

We utilized the CytoTune-iPS 2.0 Sendai reprogramming kit (ThermoFisher, A16517) to express Oct4, SOX2, KLF4 and cMyc in BJ fibroblasts. Cells were treated according to the manufacturer’s instructions three days after the second of two cycles of siRNA transfection.

### Cell culture immunofluorescence and microscopy

For immunofluorescence of human fibroblasts and hESCs cells were washed briefly with PBS, and fixed in 4% paraformaldehyde in PBS for 10 minutes. Fixed cells were washed 3 times with PBS, permeabilized with ice-cold 0.1% Triton-X in PBS for 10 minutes, and washed twice with PBS-T (PBS, 0.05% Tween-20). Samples were blocked with 3% donkey serum (Millipore Sigma, D9663) in PBS for 2 hours at room temperature and incubated overnight at 4°C using the following concentrations: anti-H3K9me3 (1:300, Abcam ab8898), anti-H3K27me3 (1:300, Active Motif, 39155), anti-SOX2 (1:200, Cell Signaling, 23064), anti-OCT4 (1:200, Abcam, ab19857), anti-PAX6 (1:200, BD Biosciences, 561462), anti-Brachyury (1:200, Cell Signaling, 81694) in 3% donkey serum in PBS. Cells were washed 3 times with PBS-T and then incubated with AlexaFluor 488- or 594-conjugated secondary antibodies raised in donkey (Thermo Fisher, See Methods) at a 1:500 dilution in 3% donkey serum in PBS.

### Total RNA isolation and sequencing

BJ fibroblasts grown in a 6-well dish were washed twice with cold PBS, followed by addition of 1 mL TRIzol reagent (ThermoFisher 15596026) and transferred to a chilled 1.5 mL tube using a cell scraper. The cells were homogenized by pipetting through a 25G needle attached to a 2 mL syringe 10 times. 200 µl of chloroform was added to the cells and vortexed for 15 seconds, followed by a room temperature incubation of 3 minutes. Phase separation of nucleic acids and protein was performed by centrifugation at 12,000 x g for 15 minutes at 4C. The upper aqueous phase containing RNA was then transferred to a fresh chilled tube. RNA was precipitated from the aqueous phase by adding an equal volume of isopropyl alcohol, followed by a 10 minute incubation at room temperature. The RNA was pelleted by centrifugation at 12,000 x g for 10 minutes at 4C. After removal of supernatant, the pellet was washed twice with 1 mL 75% ethanol, and resuspended in nuclease free water. RNA quality was verified using a high-sensitivity RNA screentape (Agilent 5067-5576). 1ug of total RNA was sent to Novogene, who assembled and sequenced bulk RNA-seq libraries. Two to three biological replicates were prepared for each condition.

### CUT&RUN experiments

CUT&RUN was performed as described in^79^, with digitonin and NaCl in wash buffers optimized for permeabilization of BJ fibroblasts and retainment of SOX2 on chromatin as performed in^80^. Next, BJ fibroblasts were detached from cell culture plates with Accutase (STEMCELL Technologies 07920), washed, bound to magnetic concanavalin A beads, and permeabilized with a dig-wash buffer (0.1% digitonin, 20 mM HEPES pH 7.5, 75 mM NaCl, 0.5 mM Spermidine, EDTA-free protease inhibitor). Bead-bound cells were incubated with a SOX2 (CST, clone D9B8N, 23064, 1:100) or H3K9me3 antibody (abcam ab8898, 1:100) at 4C overnight. The following morning, the cells were washed twice with dig-wash buffer, incubated with pA/G-MNase for an hour at 4C, and washed twice more. After chilling cells on an ice block for 5 minutes, MNase was activated for 30 minutes by addition of 2mM CaCl_2_. The reaction was stopped by adding an equal volume of 2X STOP buffer (340 mM NaCl, 20 mM EDTA, 4 mM EGTA, 0.1% Digitonin, 0.05 mg/mL glycogen, 5 mg/mL RNase A). Digested chromatin was extracted from permeabilized cells at 37C for 30 minutes, and DNA was purified by phenol-chloroform. Libraries were prepared as previously described^81^. Two to three biological replicates were prepared for each condition.

### CUT&RUN data processing

Paired-end CUT&RUN reads were mapped to the human genome build hg38 using bowtie2^82^ (v2.3.4.1) with run parameters --local --very-sensitive --no-unal --no-mixed --no-discordant -I 10 - X 1000. A quality-filtered BAM file was generated with the command samtools (v1.16.1) view -q 5 -f 2 -bS. Global scaling factors (SF) for siERH and siControl H3K9me3 data sets were calculated using the default parameters of ChIPSeqSpikeInFree^83^ (R v3.6.3). Bigwig files for visualization were generated from indexed BAM files with command bamCoverage --smoothLength 10 --normalizeUsing CPM --scaleFactor 1/SF --ignoreForNormalization chrM (deeptools 3.5.0). BED files were generated from BAMs using the bedtools command bamToBed (bedtools v2.27.1). BED files were converted to bedgraph format using the bedtools command genomecov. Peaks were called on the bedgraph files with SEACR^84^ (v1.3), using the stringent setting and selecting the top 0.01% of regions by AUC. Peak lists for downstream analyses were derived from individual replicates or the union of overlapping sites from biological triplicates, as indicated in the text and figures. Enrichment analysis of known DNA binding motifs was performed using HOMER^85^.

### Heatmaps

Heatmaps were generated with deeptools^86^ version 3.5.0. Counts matrices were generated with command computeMatrix reference-point --missingDataAsZero --referencePoint center. Figures were produced with the plotHeatmap command.

### Correlation plots

To assess genome wide similarity between hBJ siERH H3K9me3, hBJ siControl H3K9me3, and H1-hESC H3K9me3 data sets, the Pearson correlation of peak overlap was calculated using the Intervene tool^87^. In brief, peak files for individual H3K9me3 replicates were processed with the command intervene pairwise --I *H3K9me3.bed --compute fract --htype color. Visualization of Pearson correlation of fractional overlap between peak sets were generated on https://intervene.shinyapps.io/intervene.

### RNA-seq data analysis

RNA-seq reads (paired-end) were mapped to the mm10 or hg38 genome for mouse and human experiments respectively using STAR^88^ (2.7.10b). The mapped reads to genes were counted using featureCounts^89^. For counting repeat mapping reads the TEcount function from TEtranscripts^78^ was run on STAR aligned bam files using the hg38 repeatmasker gtf file annotation. The output count files were analyzed by DESeq2^90^ to detect differentially expressed genes (Benjamini adjusted p-value < 0.05, log2 fold change > 1 or log2 fold change > 1.5). Genes and repeats with fewer than 10 total mapped reads were excluded from DESeq2 and TETranscripts analysis. TPM, FPKM and RPM were calculated using an in-house R script.

### Mouse blastocyst image analysis

3D visualizations and analysis of embryos were performed using Imaris 8.2 or Imaris 9.7 software (Bitplane). The manual surface rendering module was used for cellular and nuclear segmentation, using phalloidin and DAPI signals to define cellular and nuclear boundaries, respectively. Quantifications of immunofluorescence intensity were performed by measuring the mean fluorescence intensity of voxels within the segmented object, standardized DAPI to account for light scattering.

### Cell culture image analysis

Quantification of nuclear area and signal intensity was performed using the Nuclear Morphometric Analysis (NMA) plugin for ImageJ^91^. In ImageJ saturated pixels was set to 0.35% prior to processing. NMA plugin was run using the following settings: show ellipse and use watershed options selected, minimum nuclei area=150, maximum nuclei area=6000, nuclei intensity segmentation threshold=95, nuclei insertion intensity=93%.

### Gene ontology enrichment analysis

Gene ontology enrichment analysis was performed using PANTHER^92^ (v18.0) with settings, statistical overrepresentation test: GO biological process complete, all genes in the database were used as the background reference list, and statistical significance was determined using Fisher’s Exact test with false discovery rate.

### Statistical analysis

Statistical analyses were performed in GraphPad Prism, the R software for statistical computing and with DESeq2. Pairwise comparisons were analyzed using an unpaired, two-tailed Student’s t-test. Correlation was analyzed with a two-tailed Pearson’s Correlation test. Significant overlap between gene sets was analyzed with the hypergeometric test. Differential gene expression analysis was performed using DESeq2 which calculates p-values by the Wald test and adjusted p-values corrected for multiple testing using the Banjamini and Hochberg method. Reproducibility was confirmed by independent experiments.

## Acknowledgements

R.L.M. was supported by an NIH NIDDK K01 postdoctoral fellowship DK117970-01. N.P. was supported by NIGMS (GM139970-01) and NICHD (HD102013-01A1). R35GM153180 supported work in the lab of K.S.Z.

## Author contributions

Conceptualization, R.L.M. and K.Z.; Formal analysis, A.K., B.H., H.F., K.T. and R.L.M.; Funding acquisition, K.Z; Investigation, A.K., B.H., H.F., K.T., J.Z., A.B. and R.L.M.; Methodology, A.K., B.H., R.L.M. and K.Z.; Project administration, R.L.M and K.Z.; Resources, K.Z,; Supervision, M.T-P., N.P., R.L.M. and K.Z.; Writing – Original Draft, A.K., R.L.M. and K.Z.

## Declaration of Interests

The authors declare no competing interests.

## Data availability

- RNA-seq and CUT&RUN datasets collected in this study have been deposited in the GEO of NCBI and are publicly available as of the date of publication. Accession numbers are listed in the Methods. GEO reviewer tokens for accessing the data while it remains in private status are for RNA-seq GSE268901 (wbsvkukqhzobpan), and for CUT&RUN GSE268961 (qngpsiywdvyfjyj).

## Code availability

- This paper does not report original code.
- Any additional information required to reanalyze the data reported in this paper is available from the lead contact upon request.

## Supplementary Tables

Supplementary Table 1. H3K9me3 peak coordinates in siERH human fibroblasts.

Supplementary Table 2. Fibroblast +OSKM 48 hours, SOX2 CUT&RUN peaks in siControl and siERH.

Supplementary Table 3. Fibroblast +OSKM 48 hours, siERH/siControl RNA-seq DESeq2 differential gene expression analysis results.

Supplementary Table 4. Fibroblast +OSKM 48 hours, siERH/siControl TETranscripts differential repeat expression analysis results.

Supplementary Table 5. Blastocyst immunofluorescence quantification values.

Supplementary Table 6. Blastocyst RNA-seq DESeq2 siERH/siControl differential gene expression analysis results.

Supplementary Table 7. List of genes categorized as group 1-4 by preimplantation expression from Gao et al., 2017^33^ and TPM values from this study’s siControl and siERH mouse blastocysts RNA-seq.

Supplementary Table 8. H1 hESC untreated, endoderm and ectoderm RNA-seq DESeq2 siERH/siControl differential gene expression analysis results.

Supplementary Table 9. H1 hESC untreated, endoderm, and ectoderm TETranscripts siERH/siControl differential repeat expression analysis

Supplementary Table 10. H1 hESC untreated, endoderm and ectoderm GO analysis results.

## Extended data

**Extended Data Fig. 1.**
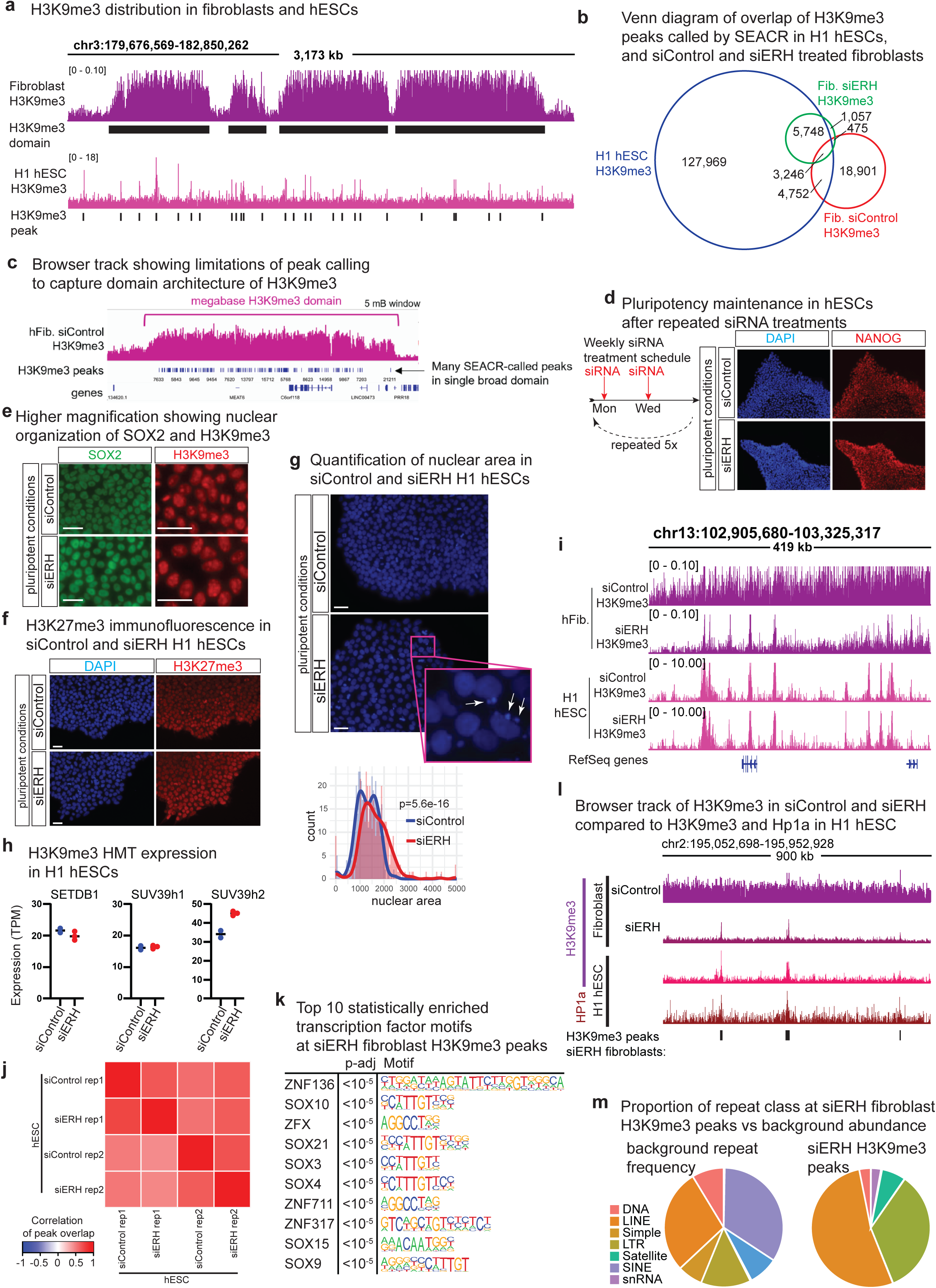
ERH depletion reverts fibroblast H3K9me3 patterns to an hESC-like state. **a.** Browser tracks comparing H3K9me3 distribution patterns in human fibroblasts compared to hESC. **b.** Venn diagram of H3K9me3 peaks called using SEACR from fibroblast siControl and siERH H3K9me3 CUT&RUN and H1 hESC ChIP-seq (GSM1003585). **c.** Browser track showing SEACR peaks called in siControl H3K9me3 domain. **d.** Nanog staining in hESCs treated for 5 weeks with siRNAs to sustain ERH depletion or control. **e.** Immunofluorescence for SOX2 (green) and H3K9me3 (red) in siControl and siERH H1 hESCs in pluripotent conditions at higher magnification to show nuclear organization. Scale bars, 50µm **f.** Immunofluorescence for DAPI (blue) and H3K27me3 (red) in siControl and siERH H1 hESCs in pluripotent conditions. Scale bars, 50µm. **g.** Nuclear area, white arrows indicate nuclear blebs. p=5.6*10^-16^, welch’s two sample t-test. Scale bars, 50µm. **h.** Expression (TPM) of H3K9me3 HMTs in siControl and siERH H1 hESCs (n=2 for siControl H1 hESCs and n=3 for siERH H1 hESCs). **i.** Browser tracks showing H3K9me3 in human fibroblasts or hESCs treated with siControl or siERH. **j.** Correlation of H3K9me3 from hESCs after siControl or siERH. **k.** Browser tracks showing H3K9me3 (GSM1003585) and HP1a (GSM1701825) in H1 hESCs and H3K9me3 in siControl and siERH human fibroblasts. **l.** Top 10 statistically enriched transcription factor binding motifs at H3K9me3 peaks in siERH treated human fibroblasts (homer findMotifs). **m.** Occurrence of repeats from RepEnrich database and repeat occurrence at sites of siERH fibroblasts H3K9me3 peaks.

**Extended Data Fig. 2.**
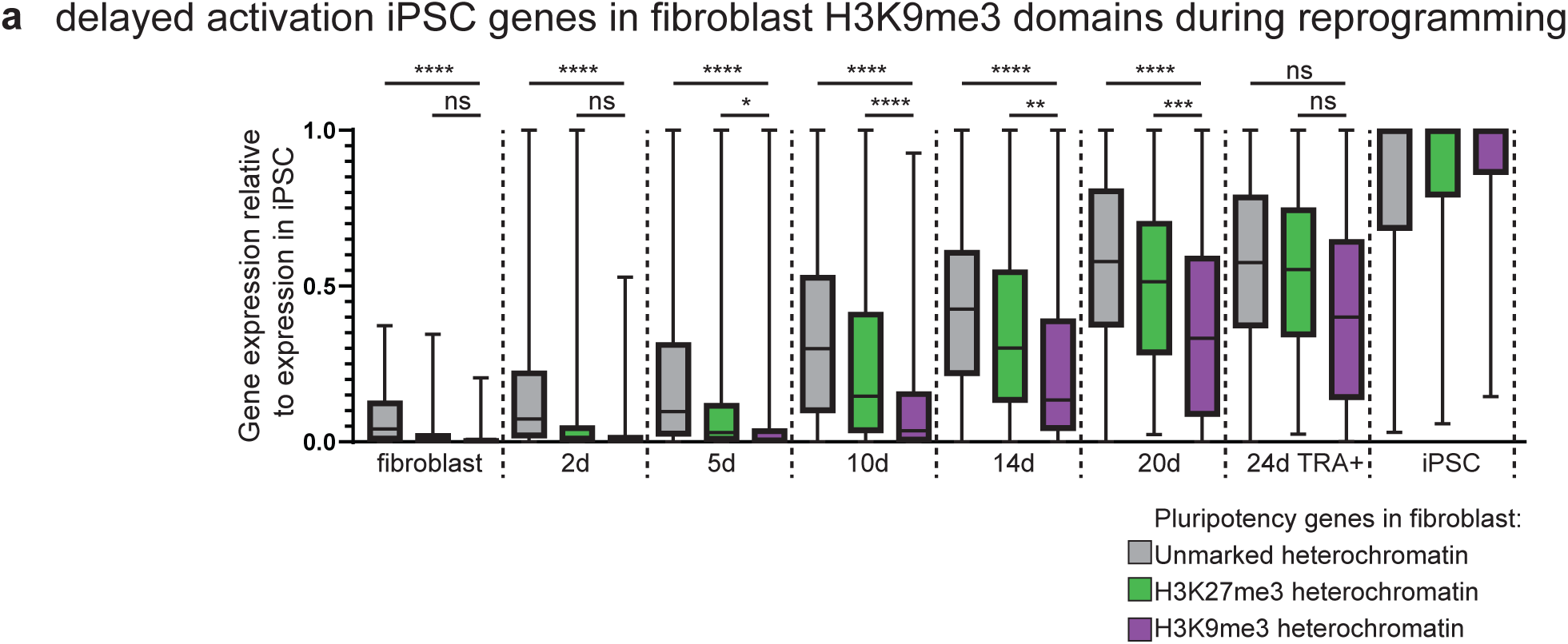
Kinetics of iPSC gene activation is effected by heterochromatin marks. **a.** Expression of genes activated during fibroblasts to iPSC reprogramming timepoints^26^ that are located in unmarked, H3K27me3 or H3K9me3 marked heterochromatin as annotated by Becker et al., 2017^5^.

**Extended Data Fig. 3.**
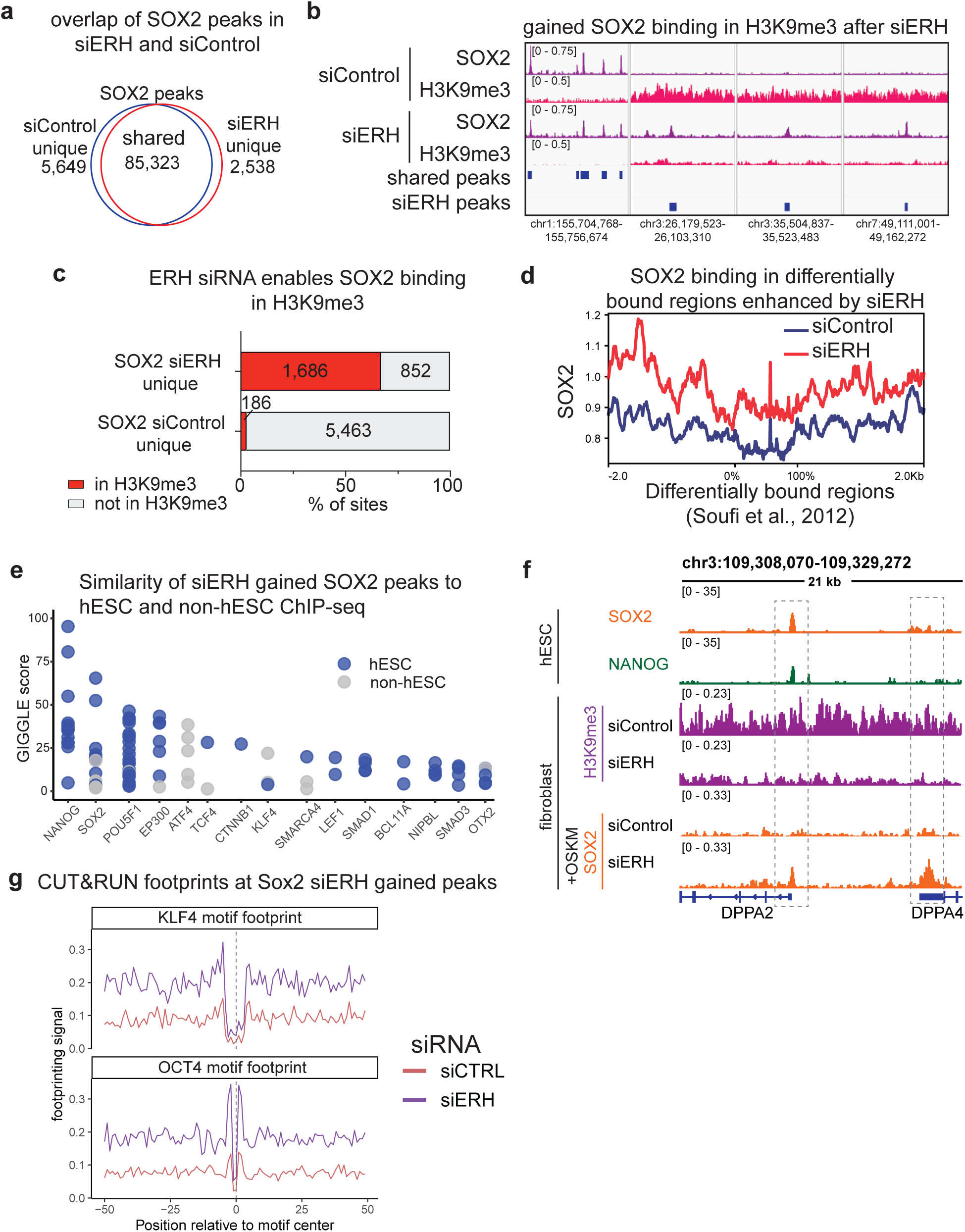
ERH depletion enhances SOX2 binding in H3K9me3 domains. **a.** Venn diagram showing overlap between SOX2 peak calls in siControl and siERH fibroblasts+OSKM. **b.** Browser track of example hESC SOX2 and NANOG peak coincident with fibroblast siERH H3K9me3 loss and SOX2 peak gain. **c.** Characterization by fibroblast H3K9me3 status of SOX2 binding sites gained in siControl and siERH fibroblasts+OSKM. **d.** SOX2 CUT&RUN average profile in siControl and siERH fibroblasts+OSKM for genome regions identified as being differentially bound by reprogramming transcription factors ^20^. **e.** Rank ordered GIGGLE enrichment scores. Each circle indicates similarity between siERH SOX2-gained sites and a ChIP-seq peak set from cistromeDB. Blue circles are peak sets profiled in hESCs, white circles are peaks from a non-embryonic origin. **f.** Browser track of H3K9me3 in siControl and siERH fibroblasts and RNA in siControl and siERH fibroblasts+OSKM at DPPA4. **g.** KLF4 and OCT4 footprint analysis at SOX2 siERH gained peaks.

**Extended Data Fig. 4.**
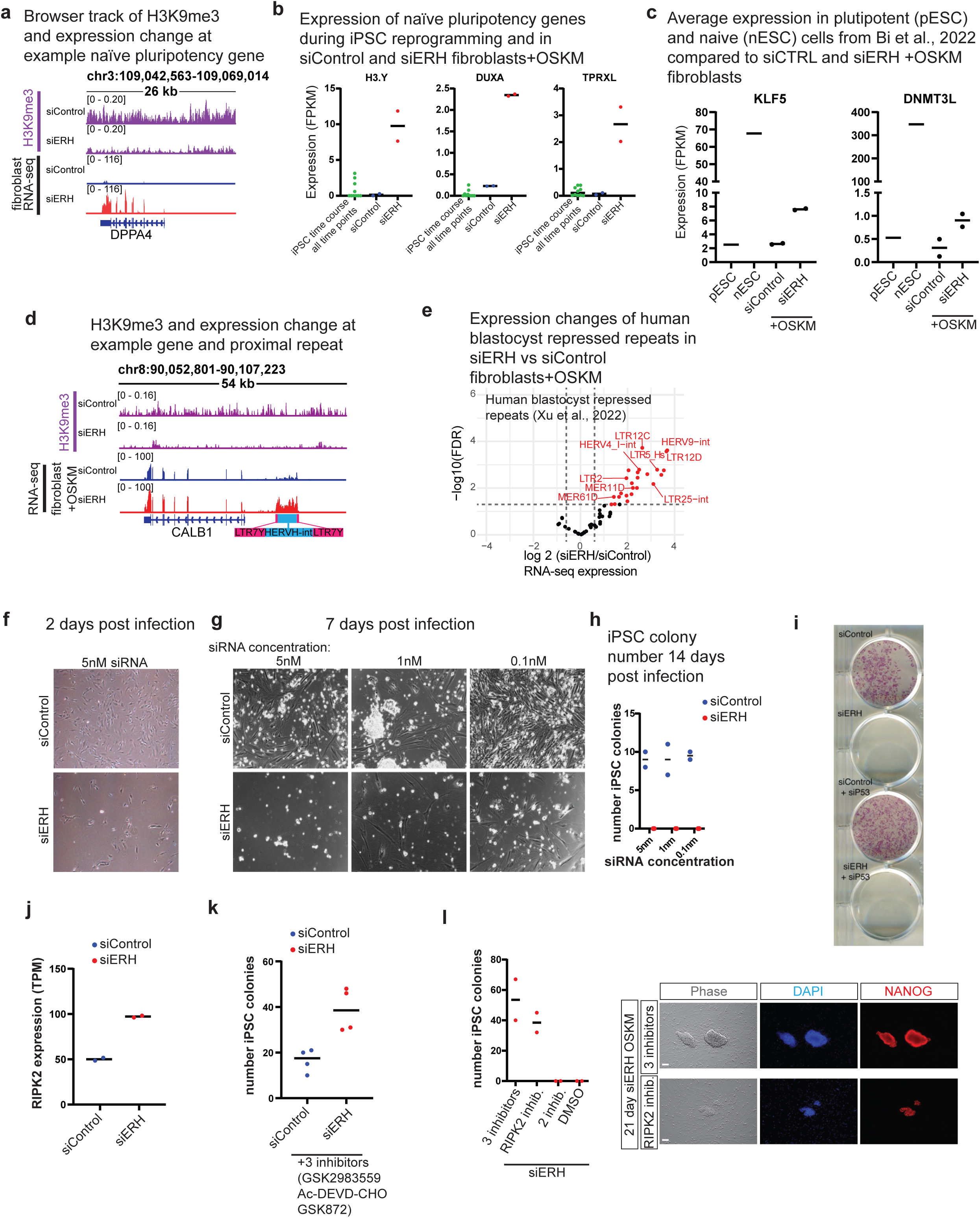
ERH depletion enhances gene activation of pluripotency. **a.** Browser track of H3K9me3 loss and gene activation of DPPA4 in siERH treated human fibroblasts+OSKM. **b.** Example 8 cell genes activated in siERH+OSKM (n=2 per condition) but not activated during eight time points of iPSC reprogramming time course^26^. **c.** Comparison of expression (FPKM) of naïve genes KLF5 and DNMT3L of siControl and siERH fibroblasts+OSKM from this study (n=2 per condition) with pluripotent and naïve human ESC data from Bi et al., 2022. **d.** Browser tracks of H3K9me3 in siControl and siERH fibroblasts and RNA in siControl and siERH fibroblasts+OSKM, at a locus of an LTR7Y-HERHV transposable element and nearby gene. **e.** Volcano plot showing differential repeat expression in siERH vs siControl fibroblast+OSKM for repeats annotated in^6^ as repressed by H3K9me3 in human blastocysts. (TETranscripts, DESeq2, padj<=0.05, log2fc>0.5). **f.** Representative phase image of siControl and siERH treated fibroblasts+OSKM 2 days after infection with OSKM sendai virus. **g.** Representative phase image showing cell confluency of siControl and siERH fibroblasts+OSKM cells 2 weeks after initial OSKM infection with titrated siRNA concentration. All treatments were plated at identical cell numbers 1 day prior to infection. **h.** Plot of iPSC colony count from siControl and siERH fibroblasts+OSKM cells 3 weeks after initial OSKM infection with titrated siRNA concentration (n=2 per condition). Each value represents the number of colonies generated in 1 well of a 24 well plate. **i.** Alkaline phosphatase staining of siControl and siERH +/- sip53 treated fibroblasts+OSKM 21 days after infection. **j.** Expression (TPM) of RIPK2 in siControl and siERH fibroblasts+OSKM 48 hours after infection (n=2 per condition). **k.** Plot of iPSC colony count 3 weeks after initial OSKM infection for siControl and siERH cells reprogrammed in media containing the inhibitors GSK2983559, Ac-DEVD-CHO and GSK872 (n=4 per condition). **l.** Plot of iPSC colony count (n=2 per condition) and imaging of siERH fibroblasts+OSKM 3 weeks after infection and treated with indicated combinations of inhibitors or DMSO control.

**Extended Data Fig. 5.**
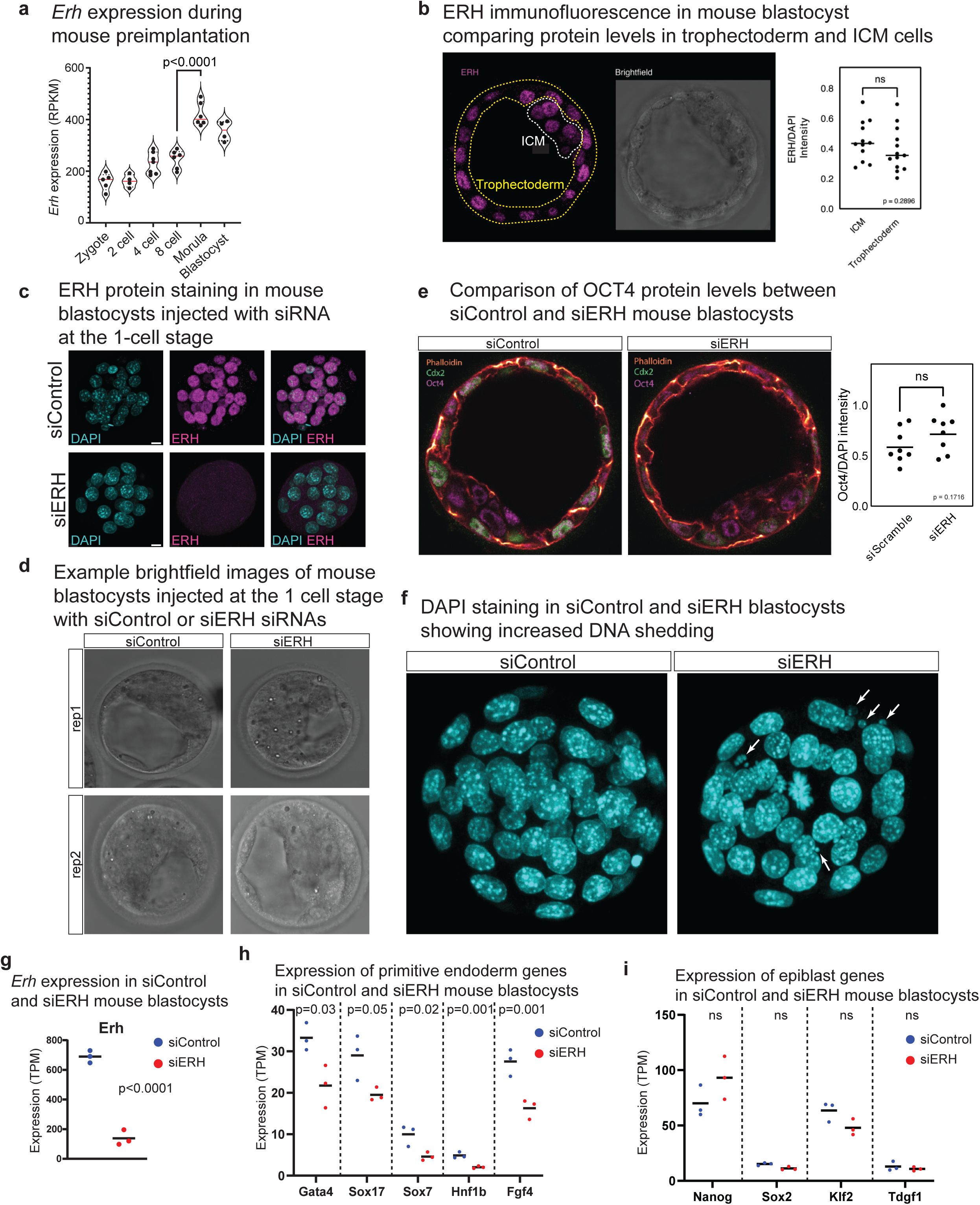
Characterization of ERH and the effects of ERH depletion in mouse blastocysts. **a.** ERH RNA expression in reads per kilobase million (RPKM) during stages of mouse pre-implantation development, RNA-seq data from^33^. unpaired two tail t-test; n=6. **b.** Fluorescent imaging of ERH (magenta) and quantification in ICM and trophectoderm cells (n=12 for ICM and n=13 for Trophectoderm, unpaired t-test, ns=not significant). **c.** ERH (magenta) and DAPI (teal) staining in mouse blastocysts injected at the 1 cell stage with siControl and siERH siRNAs showing protein depletion. Scale bars, 10µm. **d.** Representative DIC images of mouse blastocysts injected at the 1 cell stage with siControl and siERH siRNAs, two examples for each shown. **e.** Fluorescent imaging of trophectoderm marker CDX2 (green), pluripotency marker OCT4 (magenta), with Phalloidin (orange) to define cell boundary and quantification of OCT4 levels in siControl siERH blastocysts with DAPI (blue). (n=8 per condition, t-test, ns=not significant). **f.** Representative DAPI staining of siControl and siERH blastocysts. White arrows indicated DNA shedding. **g.** Expression of Erh in siControl and siERH treated mouse blastocysts by RNA-seq. (n=3 per condition, t-test, p>0.0001). **h.** TPM expression values from RNA-seq for primitive endoderm genes. (n=3 per condition, t-test) **i.** TPM expression values from RNA-seq for epiblast genes. (n=3 per condition, t-test, ns=not significant)

**Extended Data Fig. 6.**
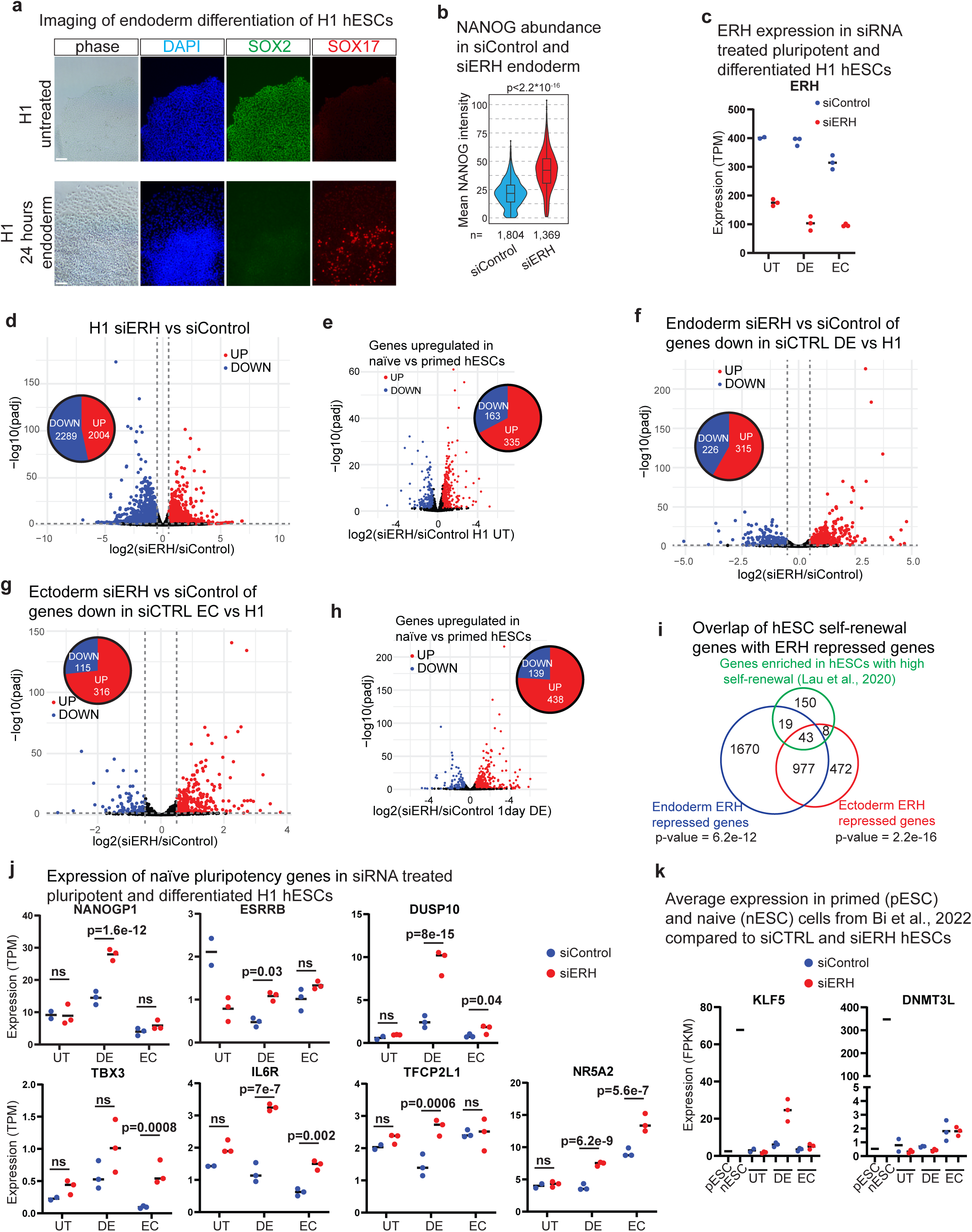
ERH is involved in silencing of pluripotency genes during differentiation. **a.** Imaging of H1 hESCs in pluripotent and 1 day endoderm differentiated showing changes in colony morphology (phase), DAPI (blue), silencing of SOX2 (green) and induction of endoderm marker SOX17 (red). Scale bars, 100µm. **b.** Quantification of nuclear NANOG immunofluorescence intensity in siControl vs siERH 1 day endoderm differentiated H1 hESCs. (unpaired t-test). **c.** ERH transcripts per million (TPM) values from RNA-seq in siControl and siERH H1 hESCs in untreated, endoderm (DE) and ectoderm (EC) differentiation conditions. **d.** Volcano plot showing differential gene expression in siERH vs siControl H1 in pluripotency conditions. Inset pie chart indicates the number of genes upregulated and down regulated in siERH vs siControl H1 (DESeq2, padj<=0.05, log2fc>0.5). **e.** Volcano plot showing differential expression of genes upregulated in naïve vs primed hESCs (List from Messmer et al., 2019^43^) in siERH vs siControl H1 in pluripotency conditions. Inset pie chart indicates the number of genes upregulated and down regulated in siERH vs siControl H1 (DESeq2, padj<=0.05, log2fc>0.5). **f.** Volcano plot showing differential gene expression in siERH vs siControl 1 day endoderm for genes down regulated in siControl DE vs siControl H1. Inset pie chart indicates the number of genes upregulated and down regulated in siERH vs siControl DE (DESeq2, padj<=0.05, log2fc>0.5). **g.** Volcano plot showing differential gene expression in siERH vs siControl 2 day ectoderm for genes down regulated in siControl EC vs siControl H1. Inset pie chart indicates the number of genes upregulated and down regulated in siERH vs siControl EC (DESeq2, padj<=0.05, log2fc>0.5). **h.** Volcano plot showing differential expression of genes upregulated in naïve vs primed hESCs (List from Messmer et al., 2019^43^) in siERH vs siControl H1 hESCs in 1 day endoderm conditions. Inset pie chart indicates the number of genes upregulated and down regulated in siERH vs siControl H1 (DESeq2, padj<=0.05, log2fc>0.5). **i.** Intersection of genes enriched in hESCs with high self-renewal^93^ with genes repressed by ERH during endoderm and ectoderm differentiation. (hypergeometric test) **j.** Graphs of naïve gene RNA-seq, transcripts per million (TPM) values from RNA-seq replicates shown (DESeq2 adjusted p-value). **k.** Graphs of key naïve gene expression in siControl and siERH H1 hESCs compared to FPKM values for naïve and primed ESCs from Bi et al., 2022. For all RNA-seq n=3, except for siCTRL UT where n=2.

**Extended Data Fig. 7.**
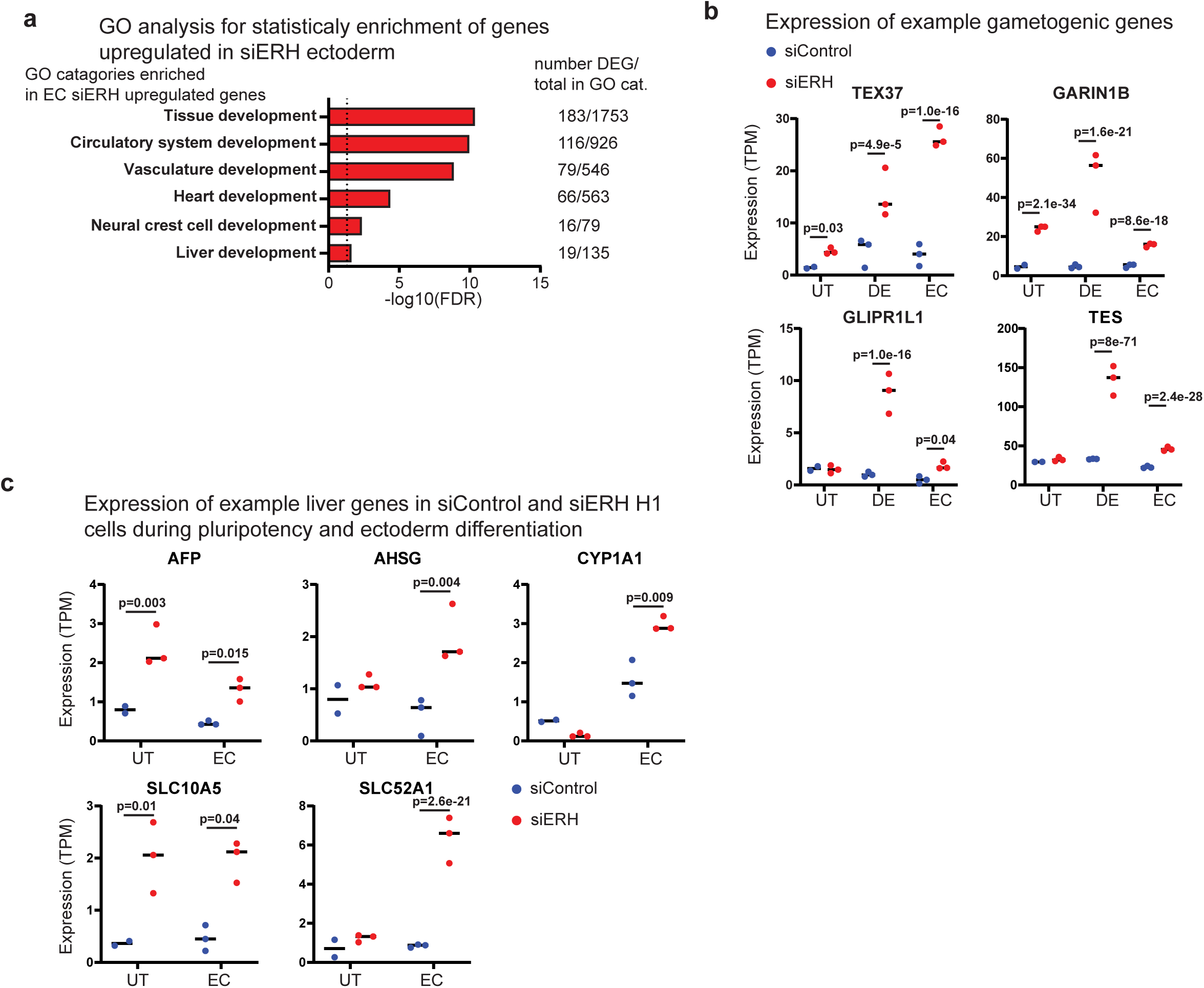
ERH represses alternative lineage gene activation during hESC differentiation. **a.** GO analysis (PantherDB, statistical overrepresentation) of genes upregulated (padj<=0.05, log2 fold change >= 0.5) in siERH vs siControl H1 hESCs 2 day ectoderm differentiation. **b.** Example meiotic and gametogenic genes activated during endoderm and ectoderm differentiation by ERH depletion, transcripts per million (TPM) values from RNA-seq replicates shown (DESeq2 adjusted p-value). **c.** Example alternative lineage genes activated during ectoderm differentiation by ERH depletion, transcripts per million (TPM) values from RNA-seq replicates shown (DESeq2 adjusted p-value). For all RNA-seq n=3, except for siCTRL UT where n=2.

**Extended Data Fig. 8.**
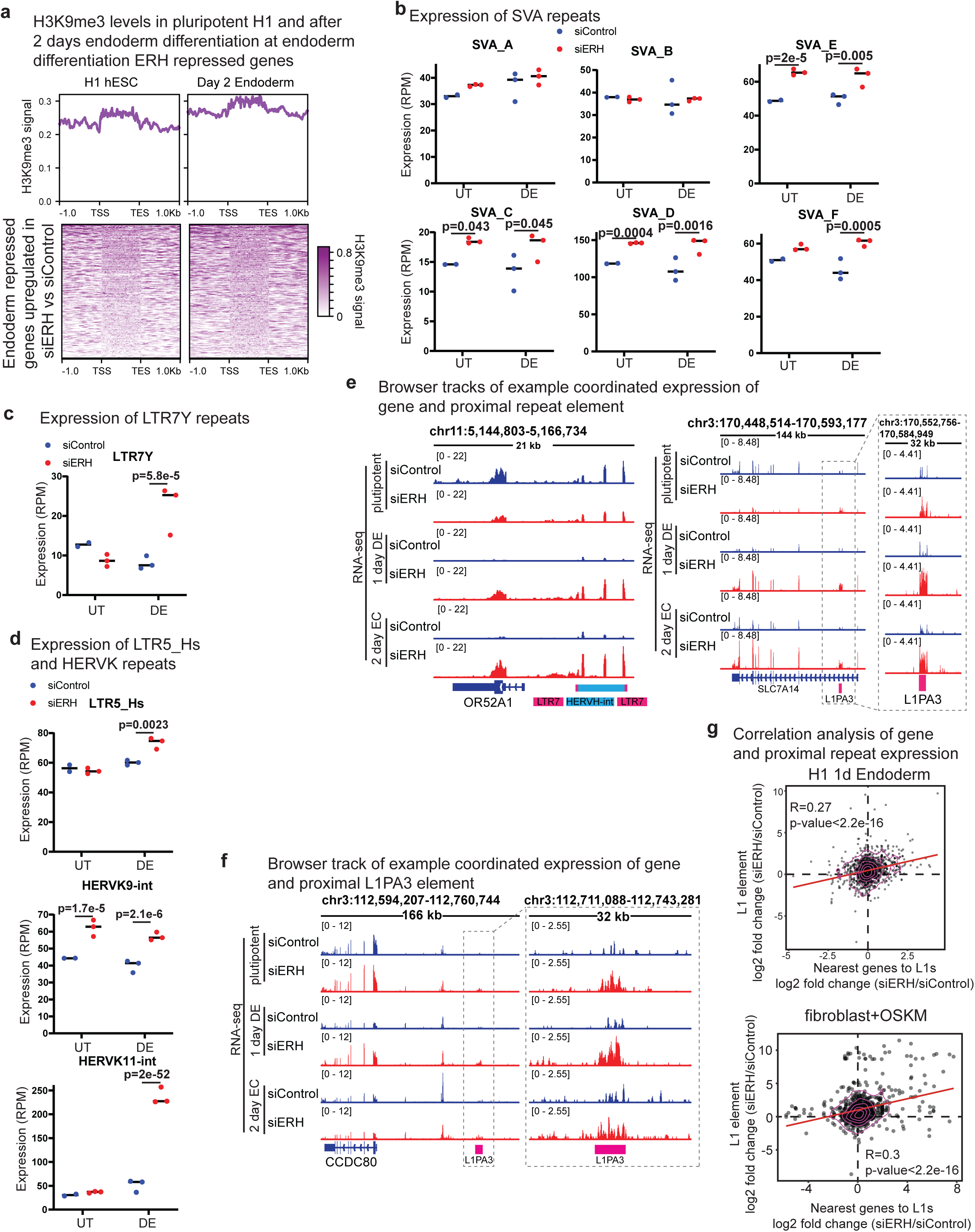
ERH represses evolutionarily young transposable elements with regulatory function. **a.** H3K9me3 data from H1 hESCs (GSM1003585) and hESC derived endoderm (GSM916057) at gene bodies +/- 1kb for upregulated (adjusted pvalue<=0.05; log2foldchange>=0.5) by ERH depletion during 1 day endoderm differentiation. **b.** SVA class element expression plots in reads per million (RPM) from RNA-seq data analyzed by TETranscripts. (DESeq2 adjusted p-value). **c.** LTR7Y expression plot from RNA-seq data analyzed by TETranscripts in untreated (UT) and endoderm (DE) differentiated H1 hESCs treated with siControl or siERH. (DESeq2 adjusted p-value). **d.** LTR5_Hs and HERVK expression plots in reads per million (RPM) from RNA-seq data analyzed by TETranscripts. (DESeq2 adjusted p-value). **e.** Browser track showing RNA-seq of example repeat element and nearby gene that is expressed in pluripotency, silenced in siControl DE and EC but expressed in siERH DE and EC conditions. **f.** Browser track showing RNA-seq of example repeat element and nearby gene that is expressed in pluripotency, silenced in siControl DE and EC but expressed in siERH DE condition. **g.** Plot of expression change of specific L1 element loci vs expression change of corresponding nearest gene within 25kb in 1 day endoderm. R, Pearson correlation. For all RNA-seq n=3, except for siCTRL UT where n=2.

**Extended Data Fig. 9.**
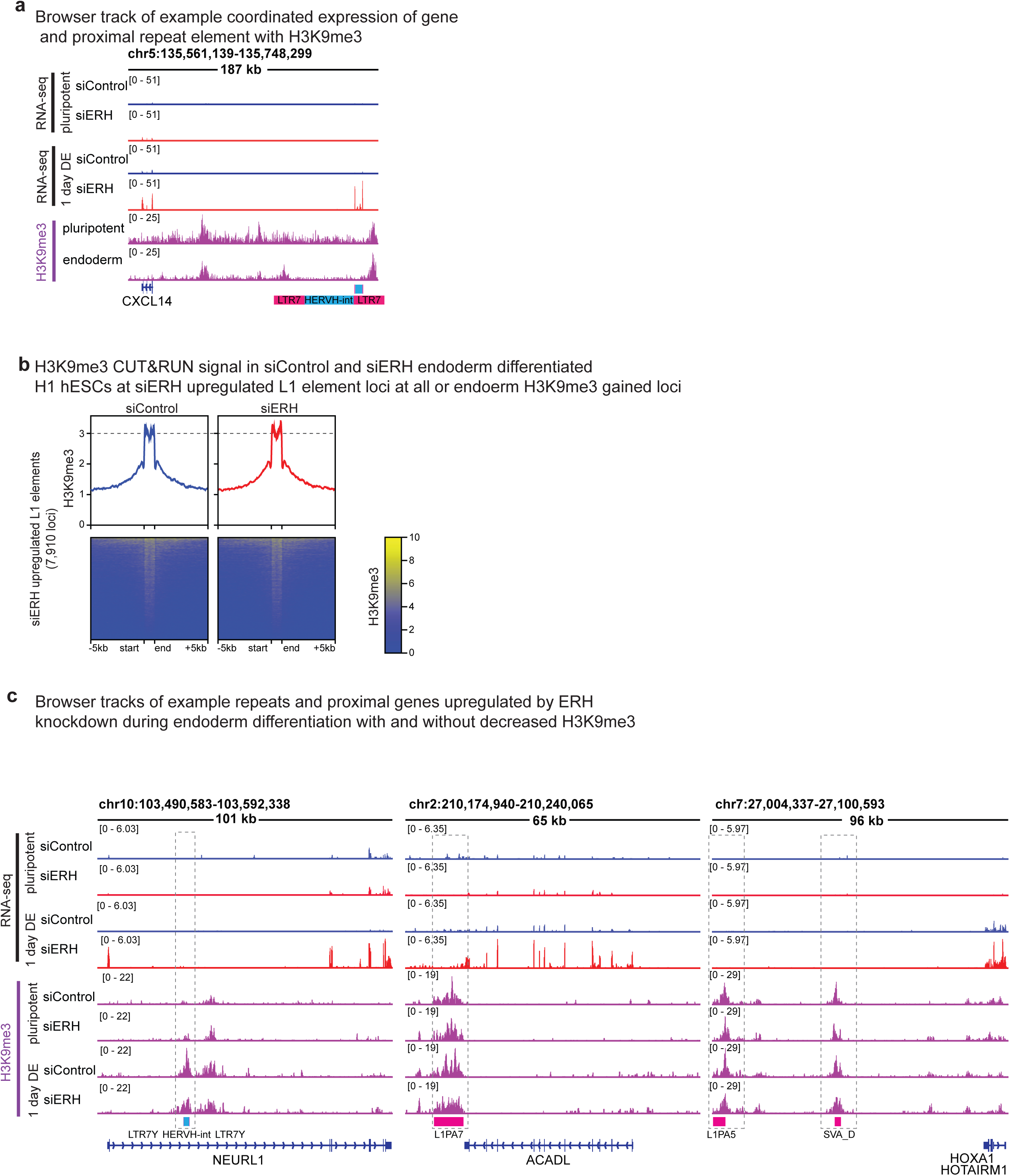
ERH dependent repression of H3K9me3 marked repeat elements. **a.** Browser track showing RNA-seq of example repeat element and nearby gene in siControl and siERH treated H1 hESCs after 1 day of endoderm differentiation and H3K9me3 tracks from H1 hESCs (GSM1003585) and hESC derived endoderm (GSM916057). **b.** Plots of H3K9me3 CUT&RUN signal at L1 elements in endoderm differentiated H1 hESCs treated with siControl or siERH. Data from loci at all L1 element loci with at least 2-fold increased RNA expression upon ERH depletion. **c.** Browser track showing example repeat element gaining H3K9me3 during endoderm differentiation in siControl and gaining less H3K9me3 in siERH. RNA-seq track showing proximal gene to repeat element being expressed in siERH endoderm.

**Extended Data Fig. 10.**
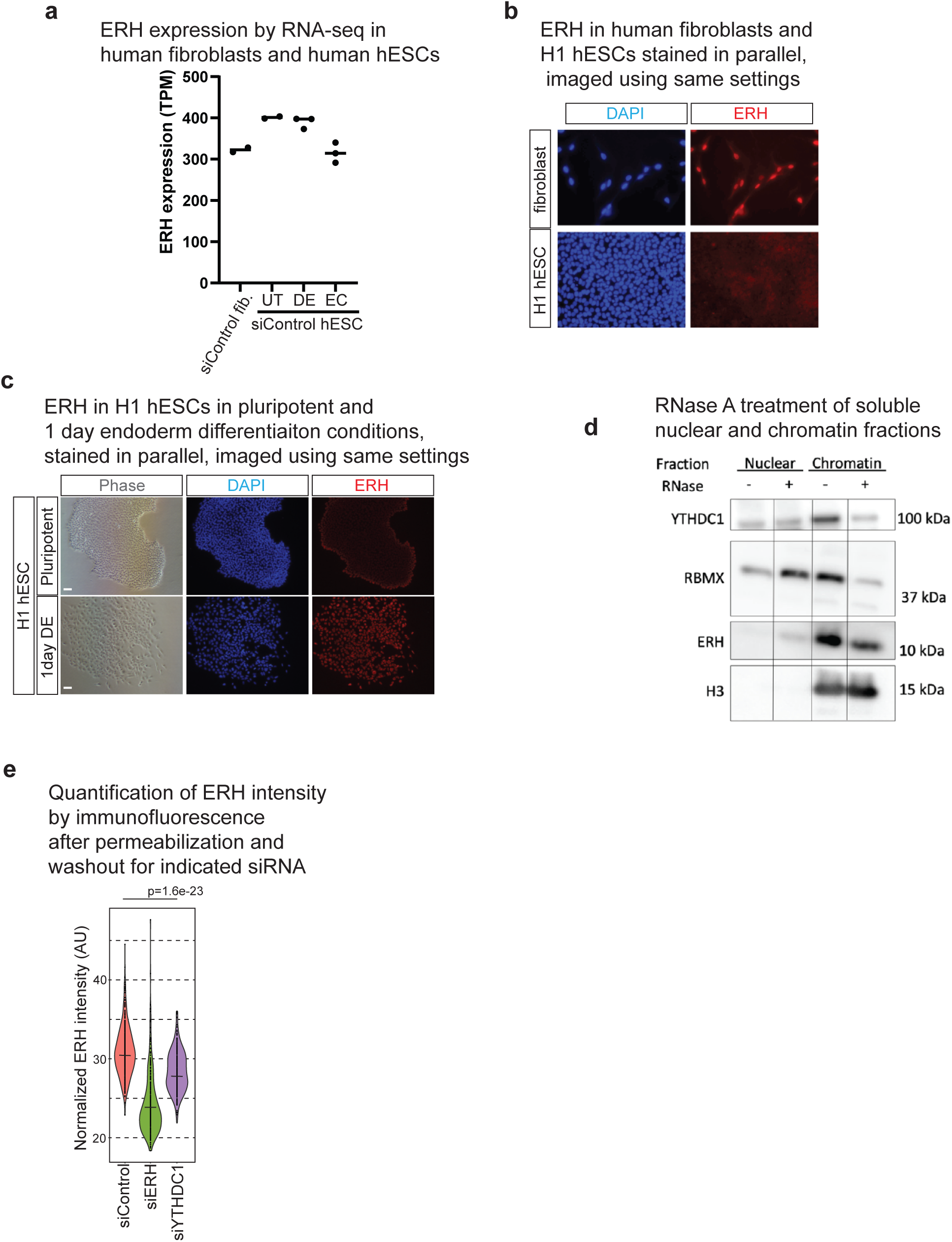
ERH protein levels are higher in differentiated cells and ERH is recruited to the chromatin by RNA and RNA binding proteins. **a.** ERH expression (TPM) from RNA-seq in siControl treated human fibroblasts, pluripotent hESCs, 1 day endoderm differentiated hESCs (DE), and 2 day ectoderm differentiated hESCs (EC). (n=2 for siControl fibroblasts and UT H1 hESCs, n=3 for siControl DE and EC H1 hESCs). **b.** ERH staining in human fibroblasts and pluripotent H1 hESCs, DAPI (blue), ERH (red). Staining and imaging performed in parallel using same conditions and microscope settings for both samples. **c.** ERH staining in pluripotent H1 hESCs and 1 day endoderm differentiated H1 hESCs, DAPI (blue), ERH (red). Scale bars, 50µm. Staining and imaging performed in parallel using same conditions and microscope settings for both samples. **d.** Western blot after RNase A treatment during soluble nuclear protein isolation. Western for ERH and YTHDC1 in nuclear and chromatin fractions with and without RNase. RBMX and H3 included as known RNA-sensitive and RNA-insensitive, chromatin-associated proteins. **e.** Quantification of immunofluorescence (IF) of nuclear ERH signal after siYTHDC1 followed by washout, in human fibroblasts.

## Notes

https://www.ncbi.nlm.nih.gov/geo/query/acc.cgi?acc=GSE268961

https://www.ncbi.nlm.nih.gov/geo/query/acc.cgi?acc=GSE268901

### Competing Interest Statement

The authors have declared no competing interest.

### Summary of Updates

This version of the manuscript has been revised with additional findings and supporting data. Additionally the manuscript has been restructured to improve clarity.

